# Balanced polymorphism at the *Pgm-1* locus of the Pompeii worm *Alvinella pompejana* and its variant adaptability is only governed by two QE mutations at linked sites

**DOI:** 10.1101/716365

**Authors:** Bioy Alexis, Le Port Anne-Sophie, Sabourin Emeline, Verheye Marie, Piccino Patrice, Faure Baptiste, Hourdez Stéphane, Mary Jean, Jollivet Didier

## Abstract

The polychaete *Alvinella pompejana* lives exclusively on the walls of deep-sea hydrothermal chimneys along the East Pacific Rise and, display specific adaptations to withstand high temperature and hypoxia associated with this highly variable habitat. Previous studies revealed the existence of a balanced polymorphism on the enzyme phosphoglucomutase associated with thermal variations where allozymes 90 and 100 exhibited different optimal activities and thermostabilities. The exploration of the mutational landscape of the phosphoglucomutase1 revealed the maintenance of four highly divergent allelic lineages encoding the three most frequent electromorphs over the worm’s geographic range. This polymorphism is only governed by two linked amino-acid replacements located in exon 3 (E155Q and E190Q). A two-niches model of selection with ‘cold’ and ‘hot’ conditions represents the most likely way for the long-term persistence of these isoforms. Using directed mutagenesis, overexpression of the three recombinant variants allowed us to test the additive effect of these two mutations on the biochemical properties of this enzyme. Results are coherent with those previously obtained from native proteins and reveal a thermodynamic trade-off between the protein thermostability and catalysis, which is likely to have maintained these functional phenotypes prior to the geographic separation of populations across the Equator, about 1.2 Mya.

## INTRODUCTION

A central goal in evolutionary biology is to understand the origin and maintenance of polymorphisms sculpted by natural selection, and more specifically how the mean phenotype of a population evolves under heterogeneous and/or changing conditions^[1]^. As a consequence, many studies have investigated the maintenance of enzyme polymorphism by selective processes for species exposed to environmental gradients such as temperature, salinity or desiccation^[2]^. A few decades ago, a series of enzymes interacting in the glycolytic cycle mostly associated with isomerase and mutase functions, such as GPI, MPI or PGM, have been shown to display isoforms that may be the subject of natural selection, leading to habitat-driven differentiation in populations according to temperature, wave action or metallic pollution^[2,3,4,5,6,7,8,9,10]^. According to Eanes^[11]^, such branch-point enzymes, which are positioned at the crossroad of metabolic pathways, are likely to be the target of natural selection as they can orientate pathway flux according to their protein variation. Amongst them, alleles encoding the enzyme phosphoglucomutase have been widely studied with the aim of testing the hypothesis of differential and/or balancing selection by looking at allele^[3,4,12]^, and heterozygote^[13]^ frequencies in populations but also to assess either the fitness of individuals carrying alleles suspected to be advantageous along latitudinal clines^[14]^ or the kinetic properties of the enzyme isoforms themselves^[6,12]^.

Because of their tremendous thermal variability due to the chaotic mixing of the cold sea water and hot fluids, hydrothermal vents represent an ideal model for testing the effect of frequent -and unpredictable-spatial and temporal changes of habitats on ‘adaptive’ enzyme polymorphisms. First, both the fragmentation and instability of the vent discharge likely promote highly dynamic meta-populations with recurrent local extinctions and associated bottlenecks^[15,16,17]^. Long-term oscillations of the heat convection beneath the ridge lead to the displacement of the hydrothermal activity along the rift, generating the emergence of new vent sites more or less closely to older ones that became extinct^[16,18]^. Second, variations of temperature, sulphide and oxygen concentrations over short periods of time (often ranging from minutes to hours^[19,20]^ are likely to affect the respiratory, nutritional and reproductive physiologies of animals living there^[21]^. Such a constant variability of the vent conditions at the individual scale represents a selective constraint that should ever promotes the emergence of highly plastic allelic isoforms or the maintenance of highly specialized alleles by favouring the heterozygous state. Finally, the hydrothermal environment is highly fragmented and heterogeneous according to the mineral composition of the oceanic crust through which the super-heated fluid is moving prior to be expulsed above the seafloor^[22]^. As a consequence, vent fields often correspond to a mosaic of edifices of different ages^[23,24]^, whose mean age and size distribution is dictated by the frequency of tectonic and volcanic events and the dynamics of the heat convection beneath oceanic ridges^[25,26,27]^. This provides distinct ecological niches for a variety of vent species able to colonise chimneys over time and thus an ecological basis for diversifying selection.

The polychaete *Alvinella pompejana*, which lives on the hottest part of the hydrothermal-vent environment^[28,29]^ can withstand temperatures up to 50°C^[30]^. This tube-dwelling worm lives on the walls of hydrothermal-vent chimneys, from a latitude of 23°N on the East Pacific Rise (EPR hereafter) to 38°S on the Pacific Antarctic Ridge (PAR)^[31]^ and has developed peculiar physiological adaptations to colonize this hostile habitat^[32,33]^. Earlier genetic studies showed that *A. pompejana* exhibits quite an unusual high level of genetic diversity^[31,34,35]^ with a non-negligible number of bi-allelic enzyme loci with equifrequent alleles, some of which displaying different thermal stabilities^[36]^. Amongst them, the enzyme phosphoglucomutase (PGM-1) possesses four distinct isoforms. Allozymes 90 and 100 have frequencies of approximately 35% and 60%, respectively in populations of the Northern EPR, the two other isoforms (112 and 78) rather rare, account for the remaining 5%. Although the frequency of allozyme 90 remained constant over the species range, Plouviez *et al.*^[35]^ however showed that allozymes 78 and 100 displayed an abrupt clinal distribution across the Equator with allozyme 78 becoming the most frequent allele in the Southern EPR. Bi-allelism was thus preserved all along the EPR despite population isolation, recurrent extinction/recolonizations and a long history of divergence across the Equatorial barrier.

In addition, significant genetic differentiation has been observed between the worm populations living in contrasted microhabitats, and especially when comparing newly formed ‘still hot’ chimneys (‘hot’ niche) to older and colder edifices (‘cold’ niche). The frequency of allele 90 was indeed positively correlated with mean temperature at the opening of *Alvinella* tubes and increased in the former habitat suggesting that this locus was under diversifying selection^[12]^. *In vitro* experiments on the enzyme stability and optimum strengthened this view and showed that allele 90 was more thermostable and more active at higher temperatures than allele 100, and thus favoured in the ‘hot’ habitat but not in the ‘cold’ one.

Although whole-length PGM sequences have now been obtained for a large panel of metazoan species, very few studies have been conducted at the population level, and most of them involved bacterial strains. While this enzyme has been extensively studied in 1970-1990’s for adaptive purposes, only few studies examined the relationship between non-synonymous changes at the gene level and the subsequent enzyme performance of the isoforms (but see^[14,37]^) for the correspondence between allelism, enzyme thermal resistance and glycogen storage in *Drosophila*). Here, we report a possible case of long-term balancing selection at an enzyme locus where alleles can be maintained by a two-niche model of selection where the proportions of the two niches greatly vary over space and time. Most of the documented cases for the long-term persistence of alleles by balancing selection, and trans-species polymorphism come from studies dealing with negative frequency-dependent selection at immune and sex-determination genes^[38,39]^. This raises questions about how a chaotic and highly fluctuating two-niche system can promote balancing selection at key branch-point enzymes.

## MATERIAL and METHODS

### Animal sampling

Specimens of *Alvinella pompejana* were collected with the ROV Victor 6000 and the deep-sea manned submersible Nautile during the cruises Phare 2002, Biospeedo 2004 and Mescal 2010 on board of the research vessel L’Atalante. Animals were sampled from targeted sites located on the North EPR (NEPR hereafter) and the South EPR (SEPR hereafter, see Fig. 1) over chimneys of different ages ranging from newly formed ‘hot’ diffusors to large black ‘smokers’, for which thermal and chemical conditions were highly contrasted^[24]^.

**Fig. 1.**
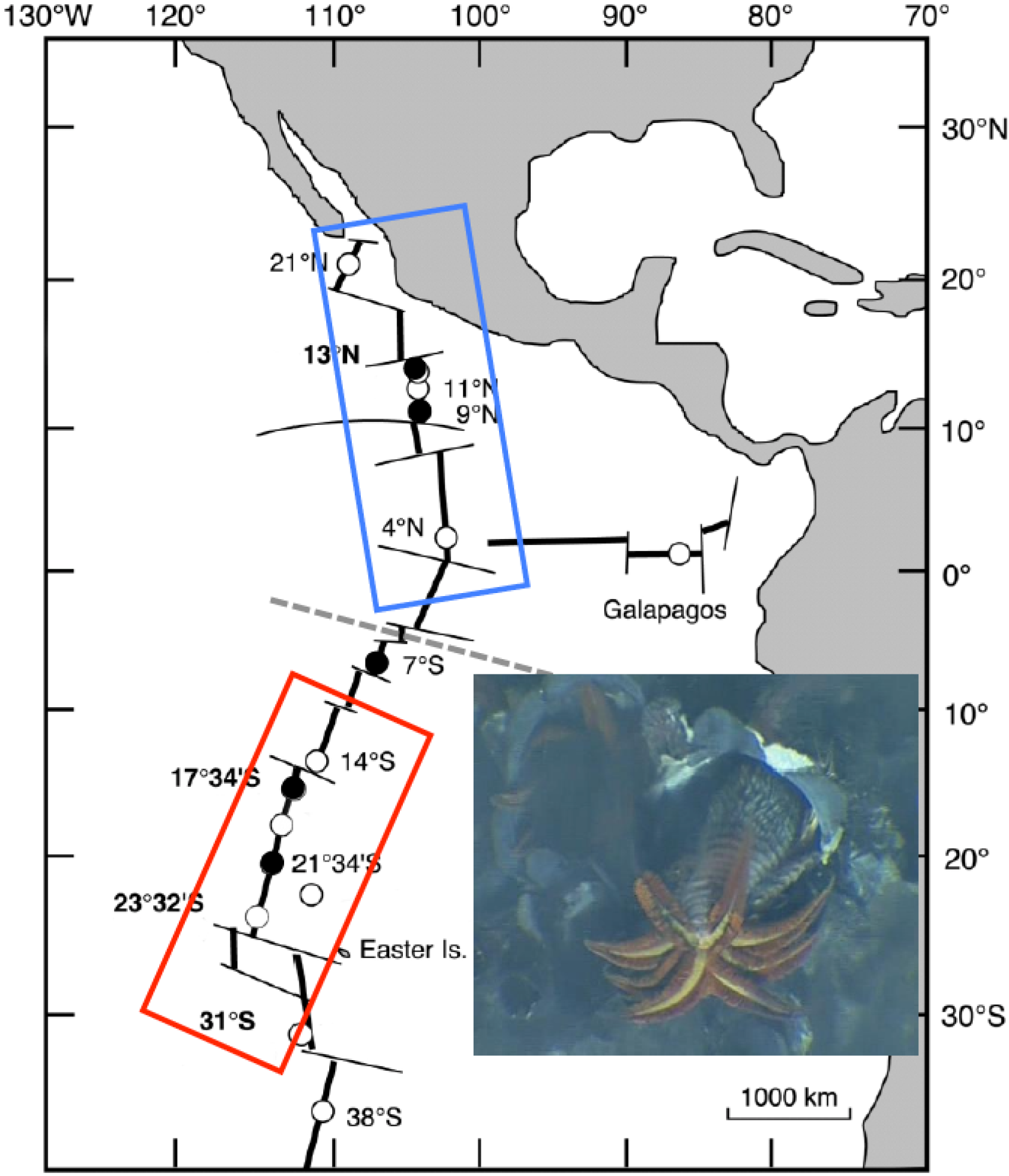
Species range of the Pompeii worm *Alvinella pompejana* along the East Pacific Rise. Dashed line indicates the presence of the Equatorial barrier to gene flow depicted by Plouviez et al. (2009, 2010[35,44]). Blue and Red boxes correspond to the northern and southern metapopulations of the worm.

In order to test an earlier hypothesis^[12]^ postulating that individuals carrying genotypes favored during the colonization of newly-formed still ‘hot’ chimneys may be counter-selected by a lower reproductive fitness under cooler conditions (i.e. trade-off between settlement ability and reproduction), we examined the relationship between PGM-1 genotypes and female’s fecundity. To this extent, the size of animals was estimated from the width at the S4 setigerous segment and sexes were determined based on the presence of either a genital pore in females or a pair of sexual tentacles in males^[40]^. Mature females collected from both sides of the Equator were dissected to estimate their fecundity per size unit and genotyped at the PGM-1 locus. For each female, the coelomic fluid containing oocytes was carefully removed and resuspended in 50 ml of a solution of borate-buffered 3% formalin in seawater. Oocytes were counted following the method previously described by Faure *et al.*^[41]^ and a one-way ANOVA was performed on size-corrected female fecundities according to the genotype using the software Jamovi (https://www.jamovi.org).

### Identification and characterization of the *AP-Pgm-1* gene

#### Sequencing of Pgm-1 cDNA using homozygous individuals

Based on allozyme genotypes, 8 homozygous individuals carrying alleles 78, 90 and 100 were selected for RNA extractions. Total RNAs were extracted with Tri-Reagent (Sigma) following the manufacturer’s instructions and a classical chloroform extraction protocol. Both the quantity and quality of RNAs were assessed with a Nanodrop ND-1000 spectrophotometer (Nanodrop Technologies, Delaware, USA). Five μg of total RNAs were reverse transcribed with a M-MLV reverse transcriptase (Promega), an anchor-oligo(dT) primer (Table S1) and random hexamers (Promega). The reverse anchor and forward nested degenerated PGM primers derived from the oyster *Crassostrea gigas* and human *Pgm-1* sequences were then used to perform the upstream amplification of cDNA fragments (see Table S1). PCR-products containing *Pgm-1* cDNA candidates were then cloned with the TOPO TA Cloning kit (Invitrogen) and, sequenced on an ABI 3130 sequencer using the BigDye v.3.1 (PerkinElmer) terminator chemistry following the manufacturer’s protocol. Sequences of clones containing the appropriate *Pgm-1* cDNA fragments were then aligned to reconstruct a series of nearly complete *AP-Pgm-1* cDNA (i.e. only lacking a small part of the 5’end of the coding sequence).

#### Sequencing the Pgm-1 gene with a series of specific exonic primers

Using gDNA, specific reverse primers (Table S1) were also used to amplify the 5’ portion of the gene by directional genome walking using PCR^[42]^. A series of specific primers were designed based on our cDNA sequences (see Table S1) to amplify both exon and intron-containing portions of the gene with gDNA from the same eight homozygous individuals. Fragments of the gene were obtained using pairs of the least distant forward and reverse primers containing a 6-bp individual identifier (barcode). PCR amplifications were performed in a 25μl PCR reaction volume that comprised 1X buffer (supplied by manufacturer), 2 mM MgCl_2_, 0.25 mM of each dNTP, 0.4 μM of each primer, 0.5 U of Taq polymerase (Thermoprime plus). The PCR profile included a first denaturation step at 94°C for 4 min followed by 30 cycles at 94°C for 30s, 60°C for 30s and 72°C for 2 min and, a final extension at 72°C for 10 min. All barcoded PCR-products were cloned following the Molecular Cloning Recapture (MCR) method developed by Bierne *et al.*^[43]^ and sequenced on an ABI 3130 sequencer with the protocol used previously. Alignments of the sequenced fragments allowed us to reconstruct a complete sequence of the *AP-Pgm-1* gene (Accession N° MN218831), its associated cDNA sequence and three native consensus cDNA for the three isoforms (Accession N° MN218832 - MN218839). The analysis of this initial cDNA alignment provided a first information on polymorphic sites between the 3 distinct alleles all along the gene (see Fig. 2).

**Fig. 2.**
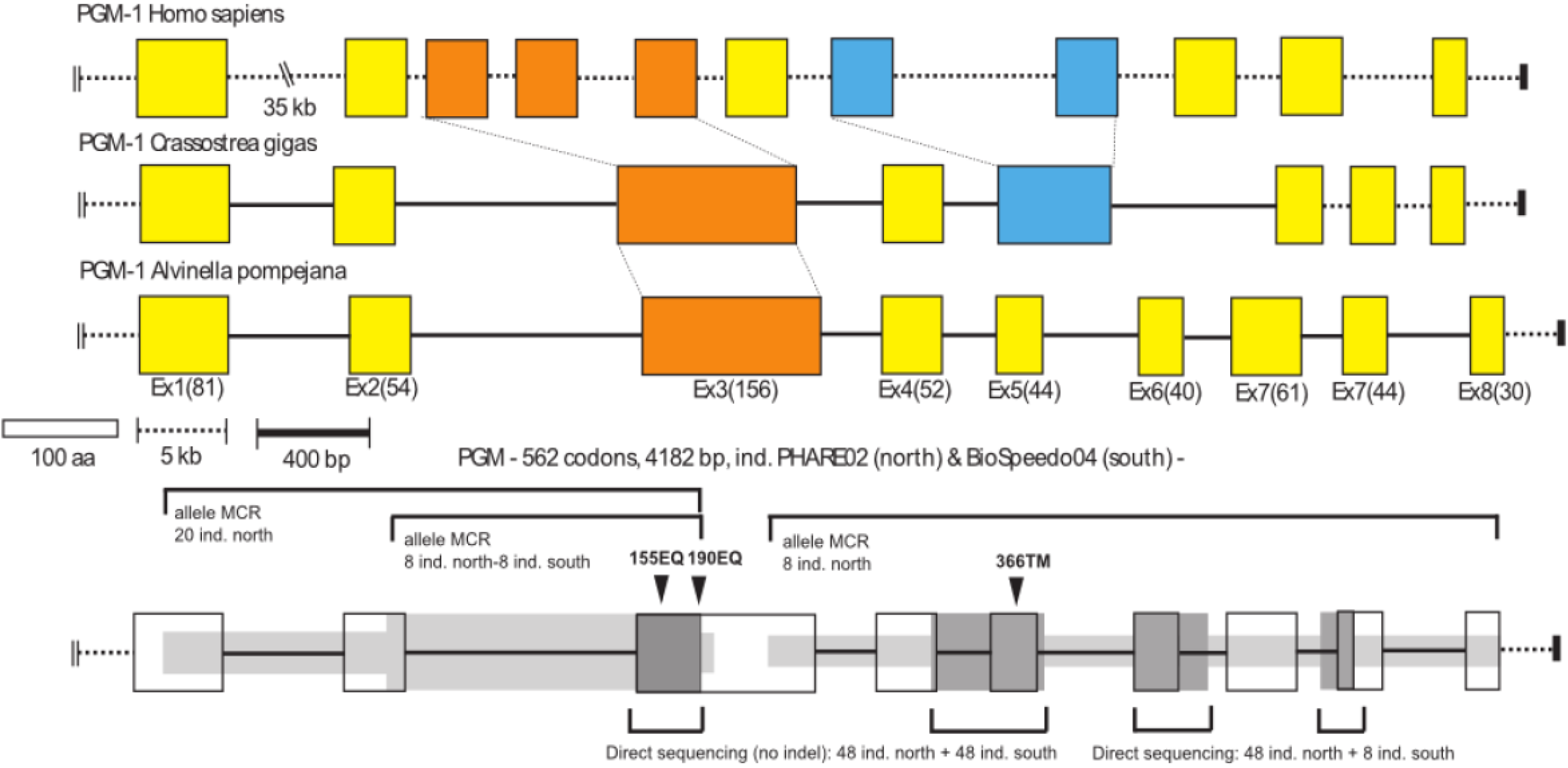
Map of the *A. pompejana Pgm-1* gene with the human (*Homo sapiens*) and the oyster (*Crassostrea gigas*) PGM as comparison. Identification of the distinct loci sequenced with the method used (Mark, Cloning, Recapture (MCR) or direct sequencing) and population origin.

### Correspondence between allozymes and non-synonymous mutations of *AP-Pgm-1*

To examine the correspondence between the only two diagnostic polymorphic non-synonymous EQ mutations found at exon 3 and allozymes 78, 90 and 100, a total of 220 individuals were genotyped on the 350bp fragment of the *Pgm-1* exon 3 containing these sites. PGM-1 allozymes were first screened for each individual by electrophoresis on 12% starch-gel at 4°C (100 V, 80 mA, 4 h) with the Tris-citrate pH 8.0 buffer system following the procedure described by Piccino^[12]^. The 350 bp exon3 fragment was then amplified by PCR on the same individuals following a gDNA extraction using a CTAB/PVP procedure described by Plouviez^[44]^. PCR amplifications were conducted using a specific primer pair (see Table S1) with a first denaturation step at 94°C for 4 min followed by 40 cycles at 94°C for 30s, 60°C for 30s and 72°C for 20s and, a final extension at 72°C for 2 min. PCR-products were first double digested on 33 individuals with enzymes *Fai I* (targeting the first substitution site) and *Bsg I* (targeting the second site) as an initial test and then sequenced without cloning on ABI 3130 automatic sequencer with the BigDye v.3.1 (PerkinElmer) terminator chemistry after an ExoSAP-IT purification.

Forward and reverse sequences were proof-read in CodonCode Aligner to check for the occurrence of single (homozygotes) or double (heterozygotes) peaks at the two polymorphic sites. The allele alignment has been deposited in Genbank (accession N° MN218918-MN219291). Linkage disequilibrium between genotypes, EE, EQ and QE and allozymes 78, 90 and 100 was tested using Linkdis^[45]^ of the software Genetix v.4.05^[46]^. The double mutation scoring among individuals allowed us to then estimate heterozygote excesses or deficiencies in populations. Departures to HDW were tested with 1000 permutations of alleles between genotypes using the same software. The exon 3 allele alignment was also used to reconstruct an allelic network using Network 4.5.1.0^[47]^, in order to examine the permeability of the equatorial barrier between populations at this locus.

### Examining the synonymous and non-synonymous changes along the AP-*Pgm-1* gene

Nucleotidic diversities were punctually assessed along the gene by combining direct sequencing and the MCR method on individuals from each side of the EPR (see Fig. 2). These regions included exon 1, exons 4 to 5, end of exon 7 and the beginning of exon 9 (Accession N° MN218840-MN218917 for exon1, Accession N° MN219292-MN219356 for exons 4 and 5). In addition, a fragment containing the whole intron 2 and the beginning of exon 3 where the two diagnostic EQ mutations are located (1110 bp) was also sequenced using the MCR method^[43]^ in order to test whether ‘hot spots’ of mutations occur around these two EQ sites but also to estimate allele divergences (Accession N° MN219357 - MN219404). In the MCR sequence sets, the number of retrieved alleles greatly varied between the different parts of the gene according to the sequencing efficiency and/or cloning success. Artifactual singletons due to the MCR method were removed by comparing the singleton rates between the MCR and direct sequencing datasets on the same fragments.

Allele diversity (Hd), nucleotide diversity (π) and its synonymous and non-synonymous components (π_S_ and π_N_), and Watterson’s theta (θ_W_) were thus examined together with deviations to neutral evolution (Tajima’D and Fu & Li’s F statistics) for both the northern and southern EPR individuals along the gene with the software DNAsp 4.10.3^[48]^ using a sliding window (size=100 and step=50). These basic genetic parameters were then compared with the critical values associated with sample size and neutral coalescent simulations implemented in the same software. Linkage disequilibrium between segregating sites and recombination among alleles were estimated by calculating the *ZnS* statistics^[49]^ together with the minimum number of recombination events (*Rm*^[50]^). The number of significant associations between linked sites was evaluated following a Fisher’s exact test and a Bonferroni correction implemented in DNAsp 4.10.3. The occurrence of recombinants was also checked using automated RDP and bootscan packages of RDP v.3.44^[51]^ and the search for hotspots in the recombination rate (4N.r) was examined along the gene (-recomb and -hotspot outputs) for both the northern and southern populations using the software Phase 2.1.1^[52]^. Genetic differentiation and allele divergence between the southern and northern parts of EPR were estimated by calculating Fst and Dxy with DNAsp 4.10.3. Genetic differentiation was tested using 1000 permutations of the sequence datasets using the randomization test developed by Hudson^[53]^. Finally, the intron 2-exon 3 alignment (1110 bp) was used to reconstruct a coalescent tree of *AP-Pgm-1* alleles in order to evaluate more specifically both the intra-locus recombination and allele divergence near the two EQ non-synonymous polymorphic sites. The evolutionary history of alleles was inferred using the Minimum Evolution method implemented in MEGA7^[54]^ using the NJ algorithm for the initial tree, pairwise deletion of ambiguous sites, and the close-neighbor-interchange (CNI) algorithm. Evolutionary distances were computed using the Maximum Composite Likelihood method.

### Coalescent simulations using models of selection

Additional simulations of a structured coalescent (N=1000 simulations with Ne=50000) were also performed using the software msms v3.2^[55]^ with an asymmetrical migration rate between two populations (pop_1->2_: 2N.m=1 and pop_2->1_: 2N.m=0.1) in a neutral way and two different hypotheses of balancing selection: (1) overdominance with SaA=500 and SAA=1, and (2) a 2-niches model of selection (4 populations and 2 habitats with SAA=1000, SaA=500 and Saa=0 in the first habitat and SAA=0, SaA=500 and Saa=1000 in the second habitat). Each set of simulations including the null hypothesis of asymmetrical migration without selection were run with two recombination rates (R=1 or 100). Population parameters including gene diversities (θ_W_, π), and Tajima’D within each deme and Fst between demes were estimated with the pylibseq 0.2.3 libraries^[56]^ using a home-made python script done by this latter author.

### Functional and structural analysis of PGM-1 recombinant isoforms

#### Plasmid construction for enzyme overexpression

Full-length *AP-Pgm* cDNA were obtained from, two homozygous individuals 100/100 (EE) and 90/90 (EQ). RT-PCR was conducted with the ClonTech SMARTer Race cDNA amplification kit following the manufacturer instructions and AP-PGMex11 reverse primer (see Table S1). These cDNAs were then used as a target to specifically amplify the complete coding sequence with primers containing cutting sites to be inserted in either Pet20b or PetDuet expression vectors (Table S1). Amplified coding sequences were double-digested with either enzymes *BamHI*/*NotI* or *AseI*/*XhoI* in a 25 μl volume containing the restriction buffer, the enzymes, and 1% BSA. The restriction products were then ligated in the appropriate expression vector after purification with a Nucleospin Gel extraction Clean up column (Macherey Nagel) and cloned into BL21DE3 *E. coli* cells amenable for IPTG induction and overexpression.

#### Directed mutagenesis

Using the full-length cDNA with the double mutation EE as a template, mutants _155_EQ and _190_EQ were produced by directed mutagenesis following the PCR protocol of Reikofski and Tao^[57]^. First amplifications were conducted in 50 μl reaction volume containing: 1X Pfu buffer containing MgCl_2_, 0.25 mM of each dNTP, 0.5 μM of each primer (petDuet and mutated primer), 0.5 U of the proof-reading *Pfu* polymerase (Promega) with 30 cycles of 94°C for 30s, 60°C for 30s and 72°C for 3 min. Secondly, the two regions of the mutated cDNA were joined following a PCR amplification without primers mixing the two previous PCR-products (1:1) under the same conditions and a final elongation step of 10 min. cDNAs containing the mutated sites _155_E->Q and _190_E->Q and the native _155_E_190_E cDNA were then sequenced on an ABI 3130 sequencer with the BigDye v.3.1 (Perkin Elmer) terminator chemistry to verify the sequences before overexpression.

#### Protein expression and purification

*E. coli* (BL21DE3) with the recombinant pETduet plasmid containing either native or mutated PGM cDNA sequences were grown into a LB medium supplied with 100 µg/mL ampicillin at 37°C until they reach an absorbance of 0.6 at 600 nm. Protein expression was induced by adding 1mM IPTG to the medium and kept at 37°C under shaking for 4 hours. Cells were then harvested by centrifugation (4°C/15 000 g/5 min), and the pellets were re-suspended in a binding buffer (20 mM Tris-HCl, pH 6.5, 500 mM NaCl, 5 mM imidazole), disrupted by French Press at 1.6 kbar. After removing cell debris by centrifugation (15 000 g/4°C/60 min), supernatants (1 μg/mL of lysate) were treated with DNAse I (Eurogentec) for 1 hour on ice. A first purification step was performed using immobilized metal affinity chromatography with a His-bind resin column (His-Bond kit, Novagen) to recover PGM variants, Protein binding with 5 mM and 60 mM imidazole and final elution of allozymes with 1 M imidazole were performed following a classical chromatography protocol (pH 6.5). The eluted fractions were concentrated using 30 kDa molecular cut-off Amicon-Ultra (Millipore^TM^). A second purification step was performed by size-exclusion chromatography (SEC) with Superdex 75 column (1 x 30) (GE Healthcare) at a flow rate of 0.5 mL/min monitored at 280 nm using a 25 mM Na_2_HPO_4_/NaH_2_PO_4_, pH 6.5. The purity of proteins was checked by SDS-PAGE stained with Coomassie brilliant blue before being kept at 4°C in an elution buffer supplemented with dithiothreitol (DTT, 10 mM) until use for enzyme assays. The protein concentrations were measured by absorption at 280 nm with the theoretical coefficient of 48,820 M^-1^.cm^-1^ as calculated using the ExPASy-ProtParam tool (http://web.expasy.org/protparam/).

#### Enzyme activity assay

PGMs activities were assayed by coupling the formation of α-D-glucose 6-phosphate (G6P) from α-D-glucose 1-phosphate (G1P) to NADPH formation using glucose 6-phosphate dehydrogenase (G6PD) as a relay enzyme. The reaction mixture contained 50 mM Tris-HCl, pH 7.4, 0.5 M MgCl_2_, 1.2 mM NADP, 0.1 µM G6PD. The recombinant PGMs were used at the following concentrations: [PGM_78_] = 0.9 µM, [PGM_90_] = 4 µM and [PGM_100_] = 0.6 µM. The concentration of the substrate (G1P) was varied from 0.2 to 60 mM to determine the kinetic constants K_m_ and V_max_ using a Lineweaver-Burk plot.

#### Thermal inactivation

The purified PGM activities were measured at 37°C at 340 nm using an UVmc^2^ spectrophotometer (Safas, Monaco) after a 30-minute incubation at challenge temperatures ranging from 5 to 60°C. Activities were then normalized as the percentage of residual activity when compared to the same sample kept in ice. A theoretical curve with the following equation was fitted to each experimental dataset using a nonlinear curve fit algorithm (Kaleidagraph 4.5.0, Synergy Software):

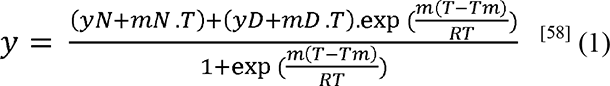

where *y* is the residual activity, *y*_N_, *m*_N_, *y*_D_, *m*_D_, the parameters characterizing the activity of the native enzyme (N) and its denatured form (D), respectively*, m* characterizing the transition between the native and the denatured forms, *R* the universal gas constant, T the absolute temperature, and *T_m_* the absolute temperature of half-denaturation, i.e. the temperature for which the activity of the enzyme is reduced by half.

#### Guanidinium chloride-induced unfolding of PGM isoforms

Unfolding of the PGM isoforms was induced by guanidinium chloride (GdmCl) in a 25 mM sodium phosphate buffer, pH 6.5, NaCl 200 mM buffer. Proteins (12 μM) were incubated with increasing concentrations of GdmCl from 0 to 5 M, 30 min at 20°C and their intrinsic fluorescence emission was determined at 324 nm under excitation at 290 nm on a Safas Xenius spectrofluorimeter (Safas, Monaco). The GdmCl concentration was determined by refractive index measurements^[59]^. Biphasic states of protein denaturation with an intermediate state (I) between native (N) and unfolded (U) states according to the following equilibrium:N↔I↔U were treated as follow: It was assumed that each transition (N↔I and I↔U) followed a two-state model of denaturation. The denatured protein fraction for each transition, *f*(I) for transition (N↔I) and *f*(II) for transition (I↔U), was determined by resolving the two following equations:

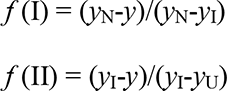

where *y*_N_, *y*_I_ and *y*_U_ are the measured fluorescence intensity respectively of the native, intermediate and unfolded state and *y* the fluorescence intensity observed at a given GdmCl concentration. The unfolded fractions *f*(I or II) data were plotted against GdmCl concentrations and theoretical curves, defined by the following equation, have been fitted on the experimental dataset using a nonlinear curve fit algorithm (Kaleidagraph 4.5.0, Synergy Software),

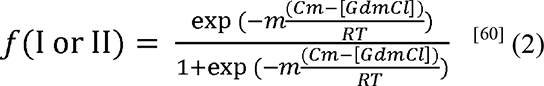

where *T* is the absolute temperature, *R* is the universal gas constant, *C*_m_ is the concentration of GdmCl at the midpoint of the transition, *m* the dependence of the Gibbs free energy of unfolding reaction (*ΔG*) on the denaturation concentration of GdmCl. Knowing *C*_m_ and *m*, standard Gibbs free energy of the unfolding reaction in absence of denaturant, *ΔG^°^_H20_*, can be calculated according to the relation:

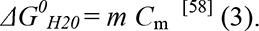

#### 3D PGM Structure Modelling

PGM 78, 90 and 100 3D protein conformations were modelled with the Modeller 9v13^[61]^, using the structure of the crystallized rabbit phosphoglucomutase with its substrate α-D-glucose 1-phosphate as a template (pdb file 1C47). This protein comprises 561 amino acids with a resolution of 2.70 Å that shares 65% sequence identity with that of *Alvinella pompejana*. One hundred models were generated for each PGM isoform and their quality was assessed using the Modeller Objective Function parameter. Finally, a structure optimization was obtained using the repair function of the FoldX software^[62]^.

## RESULTS

### Sequencing *AP-Pgm-1* cDNA from homozygous genotypes

Full-length *Pgm-1* cDNA sequences were obtained from three genotypes 100/100, three genotypes 90/90 and only two genotypes 78/78. This led to a complete cDNA sequence of 562 codons without indel between alleles encoding the three distinct allozymes (Fig. S1). The consensus protein sequence fell into the phosphoglucomutase 1 family of proteins with a blastp e-value of 0.0 (65-72% of identity over 99% of 562 residues with the sequence from the oyster *Crassostrea gigas*, and a selection of vertebrate species). Out of the 16 cloned sequences, only two non-synonymous mutations on exon 3 allowed us to discriminate the three main genotypes (100/100, 90/90 and 78/78). These polymorphic mutations corresponded to the replacement of a glutamic acid (E) by a glutamine (Q) at positions 155 and 190. Another replacement of a valine (V) by a leucine (L) at position 40 was also found in exon 1 at intermediate frequency, but this amino-acid polymorphism was not linked to a given electromorph. A phenylalanine (F) replacement by a leucine (L) was also found at position 502 in cDNA encoding allozyme 90.

### Assignment of the two EQ amino-acid replacements to allozymes in natural populations

In order to address the relationship between the two QE substitutions depicted from the cDNA sequences and allozymes, direct sequencing (and/or RFLP) were performed on a portion of exon 3 (94 codons) containing the double diagnostic mutations EQ in 220 individuals from both sides of the East Pacific Rise previously genotyped at the PGM-1 enzyme. The linkage disequilibrium between the two EQ mutations at codon positions 155 and 190, and allozymes was highly significant (Table 1) with correlation coefficients (R_ij_) greater than 70% (p-values<0.0001). This provides a very reliable correlation in which combinations QE, EQ and EE correspond to the isoforms 78, 90 and 100, respectively. The most negatively-charged allozyme 112, which is rare and always found at the heterozygous state in the northern populations was also assigned to genotype EE, suggesting that an additional replacement is occurring elsewhere in the protein. From this genotyping, groups of individuals from either the North or the South EPR did not depart significantly from the Hardy-Weinberg proportions. However, observed and expected heterozygosities were both greater in the northern population (Ho_-North_: 0.40 vs Ho_-South_: 0.29). Interestingly, allele QQ was not found in any of the populations. The frequencies of EE, EQ and QE alleles at the sampled localities are summarized in Table S2. A more thorough analysis of the North/South genetic differentiation was conducted on the 374 allelic sequences obtained by direct sequencing (see alignment in supplementary data). The resulting haplotype network (Fig. S2) shows a quasi-complete isolation of the northern and southern populations with a Fst value of 0.510 (see Table 2). Based on the 282 bp alignment, PGM90 (EQ) found in the Southern population derives directly from the northern PGM90 (EQ) by one fixed mutation and the southern PGM78 (QE) differs by 2 mutations from the northern PGM100 (EE). The haplotype network also indicated that at least three alleles sampled in the southern populations originated from the northern populations, suggesting that the barrier to gene flow is not completely sealed.

**Table 1.** Linkage disequilibrium between the combination of the two diagnostic mutations EQ and PGM-1 allozymes.

| Mutation | $D_{ij}$ | $R_{ij}$ | $K\chi^2$ | p-value |
| --- | --- | --- | --- | --- |
| EE-100 | 0.274 | 0.908 | 87.2 | 0.0001*** |
| EQ-90 | 0.115 | 0.725 | 55.7 | 0.0001*** |
| QE-78 | 0.357 | 0.907 | 87.2 | 0.0001*** |
| EE-112 | 0.013 | 0.230 | 5.6 | 0.0178* |

**Table 2.**
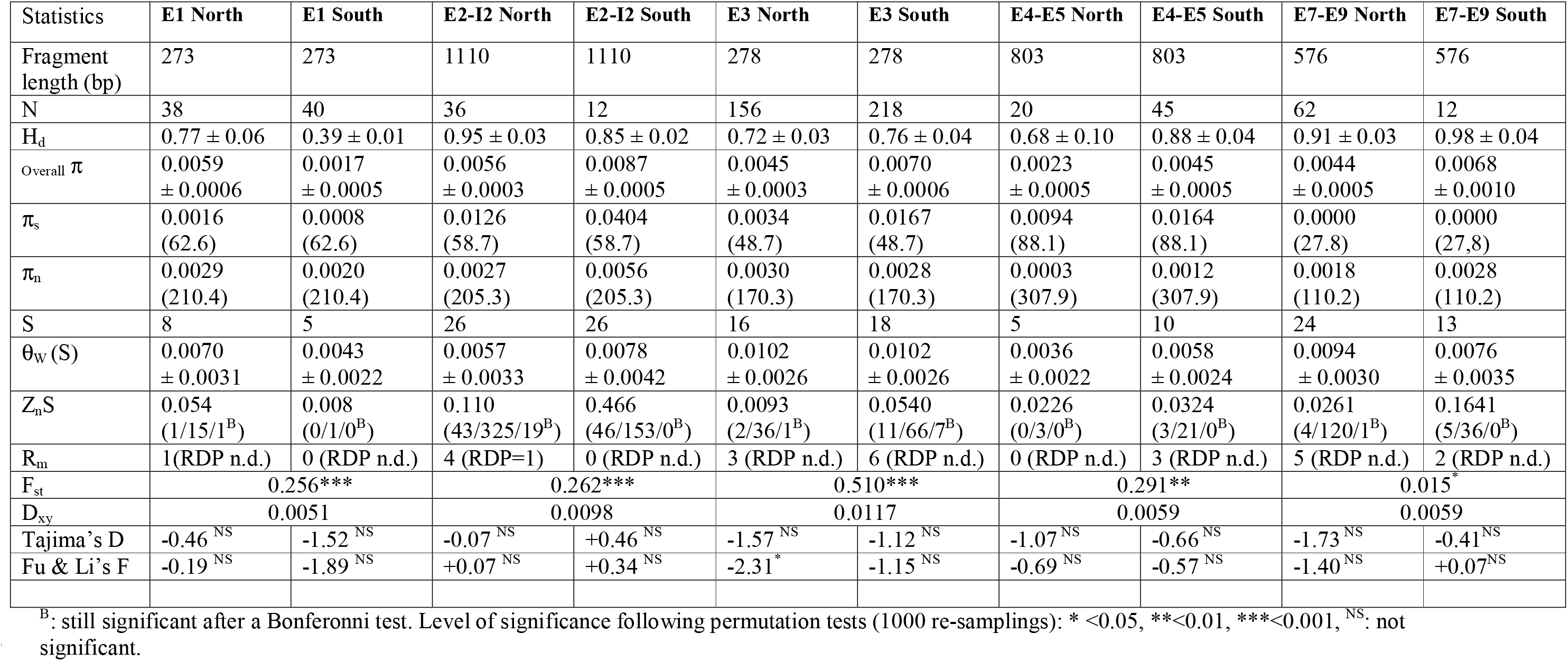
Gene diversities, population parameters and neutrality tests along the *Pgm-1* gene for *A. pompejana* populations of the South and North EPR. N and S represent the number of sequences and the number of segregating sites used, respectively. Linkage disequilibria between sites were only estimated between informative sites only: numbers in brackets correspond to the number of significant exact Fisher tests, total number of comparisons and numbers of tests still significant after the Bonferonni correction, respectively. (RDP n.d.): recombinant not detected using automated RDP and bootscan packages of RDP v.3.44. Values in brackets below π_S_ and π_N_ (Jukes & Cantor estimates) are the numbers of synonymous and non-synonymous sites in coding regions, respectively. All genetic datasets obtained using the MCR method were corrected for artifactual/somatic singletons.

### Cryptic amino-acid variation along the *AP-Pgm-1* gene

The full sequence of the *AP-Pgm-1* gene with the location of polymorphic codons and primers are shown in Fig. S1. The total length of the nucleotidic sequence is 4372 bp. The gene is subdivided into 9 exons and 8 introns which length ranges from 155 to 848 bp. The coding sequence of 1686 bp (562 codons) has an overall GC content of 43.5% (compared to only 29.3% in the intronic regions). When compared to human and oyster *Pgm-1* genes^[63,64]^, the largest *AP-Pgm-1* exon, comprising 156 codons (other exons vary from 40 to 81 codons), corresponds to the fusion of exons 3, 4, and 5 of the human *Pgm-1*. This fusion is shared with the oyster *C. gigas*, suggesting that annelids and mollusks are sharing the same gene architecture (Fig. 2).

Besides the two QE changes affecting the net charge of the protein in exon 3, other, less common, cryptic amino-acid replacements were found along several regions of the *AP-Pgm-1* CDS. This allowed us to estimate gene diversities and the south/north divergence over an overall portion of about 3 kb (two thirds of the gene, see Table 2). Gene diversities were almost constant over the *AP-Pgm-1* gene but allele divergence increases locally in the vicinity of the two segregating EQ sites (Table 2). Looking more closely at the site variation along the gene using a sliding window on our set of sequenced fragments indicated that gene diversity is also slightly higher in exon3 where the two QE substitutions are found with values almost identical to those depicted in intron2 (Fig. 3). This slight increase corresponded to peaks of positive Tajima’s D values, which raised up to +0.5 at the beginning of exon3, suggesting that the presence of the two linked non-synonymous mutations may be associated with a hotspot of gene diversity. Observed genetic diversities as estimated from θ_W_ and π were however not significantly greater than expected from neutral coalescent simulations for both the southern and northern populations over all the investigated *Pgm1* fragments (Fig.3, Table 2). Together with the two QE variant sites, the genotyping of exon 1 also confirmed the occurrence of a trans-equatorial V40L substitution found at a frequency of 0.15 restricted to the southern EQ allele (PGM90) and one of the two northern allelic lineages, irrespective of the mutations EE (PGM100) and EQ (PGM90). The direct sequencing of the two other genic regions located either between exons 4 and 5 and between exons 6 and 8 did not show any additional diagnostic amino-acid changes between the 3 allelic lineages EE, EQ and QE. By contrast, several synonymous changes and indels appear to segregate between different allelic lineages along the gene (see sequence alignments provided as supplementary data for exons 1, 3, 4, 5, 7 and introns 2, 6, and 7).

**Fig. 3.**
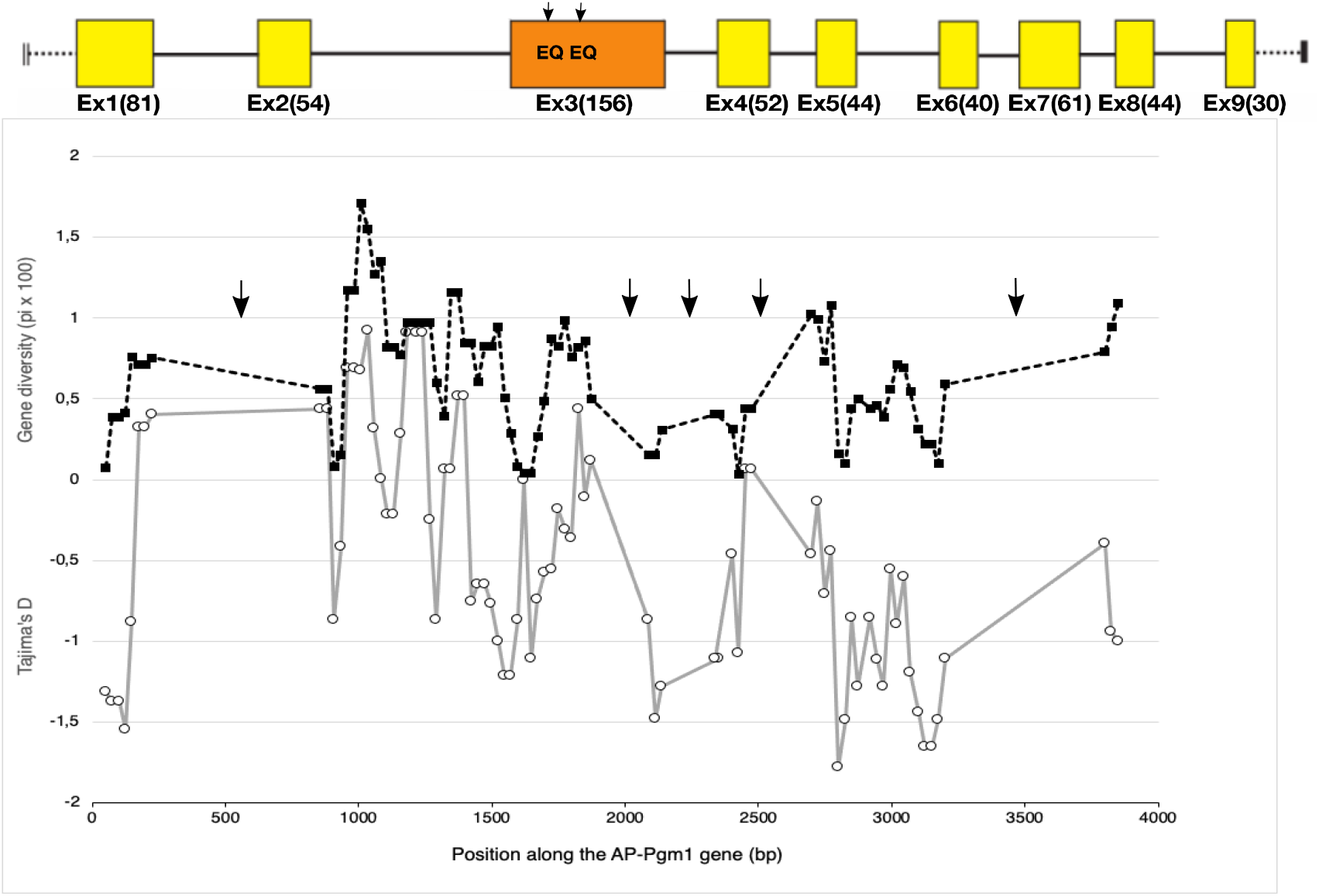
Evolution of gene diversity (π) and the statistic Tajima’s D along the *AP-Pgm-1* gene using a sliding window of 100 bp size and a step of 25 bp. The analysis includes exonic and intronic fragments for which the sequence polymorphism has been documented. Arrows indicate the portions of the gene for which there are no genetic dataset.

### Estimating allele divergence and linkage disequilibria between segregating sites

To examine more specifically allele divergence and linkage disequilibria between segregating sites within allelic lineages, a sequencing of recaptured alleles was targeted on the longest region of the *AP-Pgm-1* gene (1110 bp). This region containing intron 2 and the two allozyme-diagnostic sites EQ on exon 3 was thus genotyped from a subset of individuals. The sequencing of 48 alleles highlighted the presence of a high level of synonymous polymorphism with a strong linkage disequilibrium between these sites (Table 2), and two diagnostic indels in intron 2 (insertions referred to as A and B following their order in the intron). These segregating sites and indels allowed us to determine 4 divergent allelic lineages with a few recombinants between them. These alleles were split between the northern and southern populations. In the southern population, the two allelic lineages L1 and L2 refer to the allozyme-diagnostic double mutation QE and EQ, whereas allelic lineages L3 and L4 refer to EQ and a mixture of EQ and EE in the northern population (Fig. 4). At least, 9 and 11 synonymous mutations were fixed in intron 2 between allelic lineages L1 and L2, on one hand, and L3 and L4, in the other hand.

**Fig. 4.**
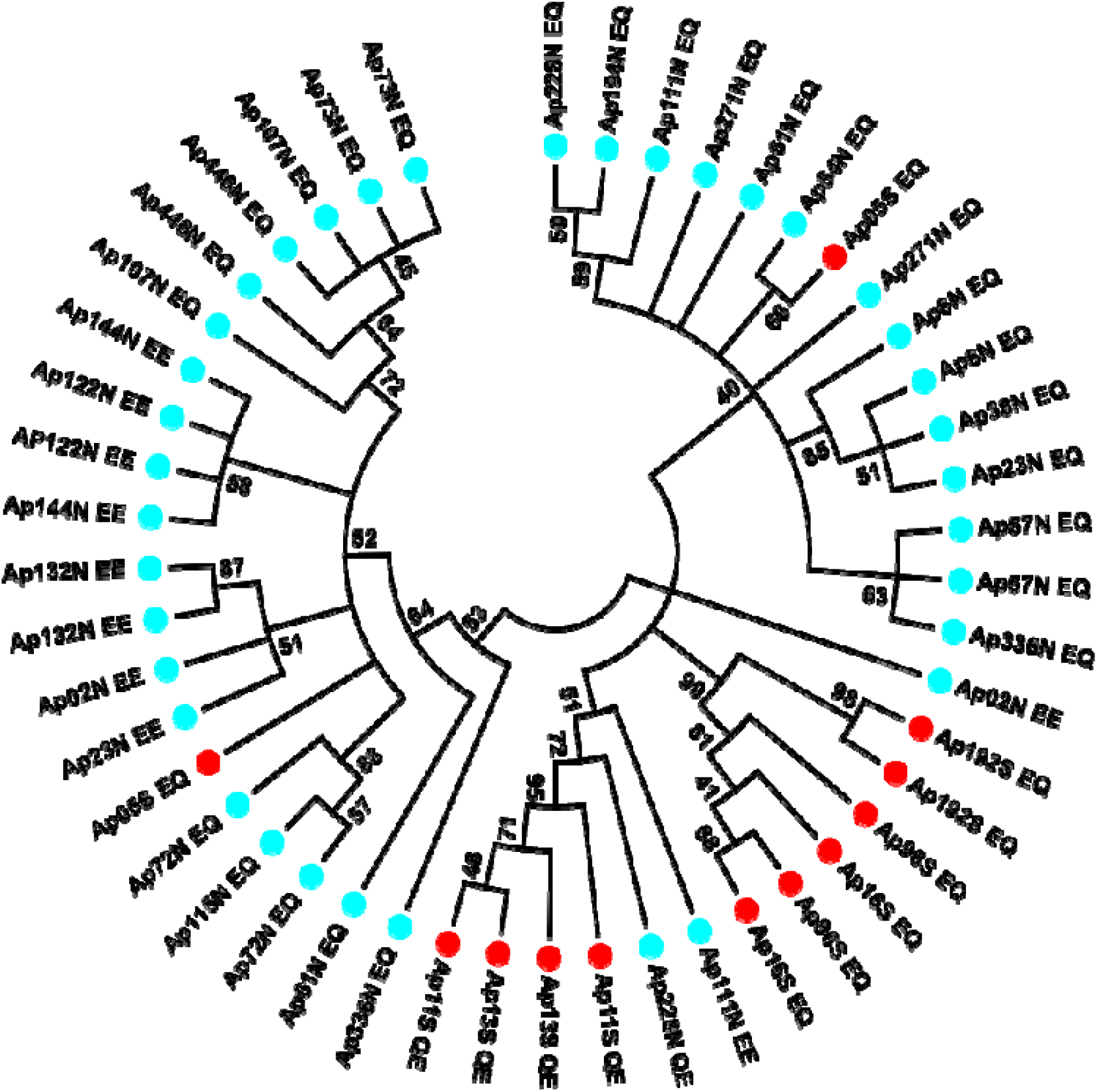
Minimum evolution tree obtained from evolutionary distances computed with the Maximum Composite Likelihood method with MEGA7 on 48 sequences from individuals of the North and South EPR locations using the Mark-Cloning-Recapture (MCR) of the *Pgm-1* introns 2 and exon 3 (1110 bp). The sequences corresponding to the PGM 78, 90 and 100 are respectively identified by the letters QE, EQ and EE traducing the polymorphism at positions 155 and 190, with the colours blue and red corresponding respectively to the individuals from the north and the south.

In the Southern population, allele L1 is typified by no insertion (QE, no_A, no_B) when compared to allele L2 (EQ and A, B) with a strong linkage disequilibrium between nearly all segregating sites (no recombination, see Table 2). It is however worth noting that one individual presently sampled at 7°25S originated from the northern populations with a L3L4 signature.

In the Northern population, the two divergent lineages L3 and L4 also display linked sites with either the A indel (L1) or the B indel (L2) but these two lineages are not completely associated with the double mutations EE and EQ. Alleles EE were only found in one of the two lineages and one recombinant between L3 and L4, suggesting that these two lineages have recombined once (Fig 4; Table 2). Alternatively, allele EE could derive from one of the two lineages.

To estimate the recombination rate, we examined the distribution of the *Rho* parameter 4N.r with Phase 2.1.1 over a greater proportion of the gene (1-2860 bp) using segregating sites (n=53) shared between northern and southern individuals that were successfully sequenced for all exon-intron fragments of the *AP-Pgm-1* gene (N=20). Results from the Phase -recomb and -hotspot outputs clearly indicated that the recombination rate between alleles remains extremely low all along the gene (average local *Rho*= 0.033 and 0.008 for the northern and southern populations, respectively, which further increased to nearly 2 in the southern population at the end of the gene near the position 2140. This study therefore indicated that the 4 allelic lineages greatly diverge one to each other in the vicinity of the double mutation characterizing allozymes, with divergence even greater between allelic sequences of the same population (0.7-1%) than those of the two geographic regions investigated (0.9%).

To test whether the *AP-Pgm1* genetic patterns may be maintained by selection, population parameters of both the northern and southern populations were simulated using a msms structured coalescent with and without selection. Simulations indicated that a low asymmetrical migration across the equatorial barrier with low or no recombination does not explain by itself the observed patterns of genetic diversities found for the *AP-Pgm-1* gene (Table S3). Simulated Fst values were around 0.8 and asymmetrical deme diversities were at least two times smaller than the observed ones (π = θ_W_ =3) with and without recombination. In this context, Tajima’s D was close to zero within each deme as observed but highly positive (+2.75) for the overall population when the observed one was also close to zero. Introducing selection led to a better fit of simulated parameters to the observed ones. Simulations with overdominance and low recombination led to a slight decrease of Fst values (0.7) between demes, an increase of the within-deme genetic diversities close to the observed ones but also produced greater positive Tajima’s D (+1.3 for each deme). Our best fit to values observed in the worm’s populations was obtained with the two-niches model simulations (Fst=0.45, converging nucleotidic diversities (θ_W_=17->13) and Tajima’D (+0.8->+0.4) estimates within and between demes). These simulated values were even closest to those estimated in the vicinity of the two EQ sites (intron2, see Tables 2, S3).

### Conformational stability, thermal inactivation and kinetics of the mutated isoforms

The obtention of full-length *AP-Pgm-1* cDNAs allowed us to examine the direct effect of the two QE substitutions on the thermal stability and efficiency of the PGM-1 enzyme using *in vitro* directed mutagenesis. To determine the conformational stability of the three recombinant isoforms of the PGM-1, their guanidinium chloride (GdmCl)-induced unfolding was studied. Variations of fluorescence with increasing concentrations of GdmCl were biphasic (Fig S3) suggesting that the protein follows a three-state model of denaturation. For each transition, the unfolded fraction of protein (*f_u_*) was determined (Fig S4) and Gibbs free energy change associated with each transition calculated (Table 3). For the two transitions, the PGM90 (EQ) appears more stable than the two other isoforms. PGM78 (QE) appears more stable than PGM100 for the first transition (*ΔG^°^_H2O_* = 8.0 ± 0.46 kJ.mol^-1^ *vs. ΔG^°^_H2O_* = 6.06 ± 0.81 kJ.mol^-1^ respectively), but not for the second transition (*ΔG^°^_H2O_* = 15.43 ± 0.93 kJ.mol^-1^ *vs. ΔG^°^_H2O_* = 15.13 ± 0.98 kJ.mol^-1^ respectively). The *T*_m_ values obtained from the theoretical curve fitted on the thermal inactivation experimental data (inset Fig. 5) are very similar for PGM78 (46.5±1.7°C) and PGM100 (44.0±0.1°C), but markedly higher for PGM90 (50.9±0.7°C).

**Fig. 5.**
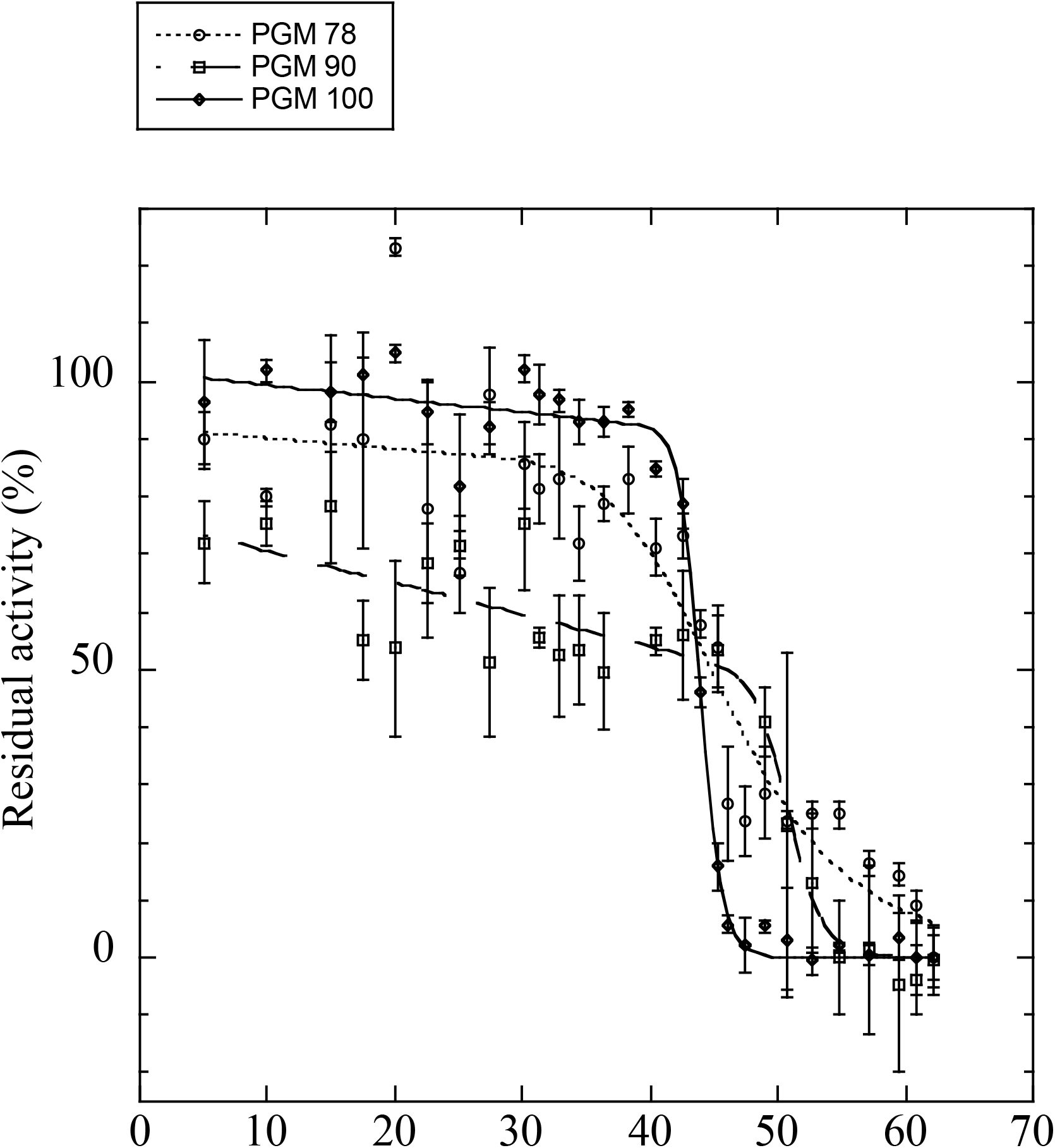
Residual enzyme activities after 30 min of incubation at different temperatures for the overexpressed isoforms of PGM 78 (QE), 90 (EQ) and 100 (EE). T_m_ values are shown in Table 4.

**Table 3.** Conformational and temperature stability of the three overexpressed variants (PGM78, PGM90, and PGM100). *Cm* et *m* values estimated from the variation of protein fluorescence in presence of an increasing concentration of GdmHCl (values for each of the two transitions). Estimation of the free enthalpy of the unfolding reaction in absence of chaotropic agent for each of the two transition states. *Tm*: values of the temperature at which we reach 50% of non-reversible inactivation after a 30-minute exposure *K^app^_m_* and *K_cat_* are kinetic parameters corresponding to the apparent Michaelis-Menten constant for glucose-1-phosphate, and the catalytic constant, respectively. The ratio of these two values corresponds to the specific activity.

|  | PGM 78 |  | PGM 90 |  | PGM 100 |  |
| --- | --- | --- | --- | --- | --- | --- |
|  | 1<sup>st</sup> transition | 2<sup>nd</sup> transition | 1<sup>st</sup> transition | 2<sup>nd</sup> transition | 1<sup>st</sup> transition | 2<sup>nd</sup> transition |
| $C_m$ (M) | $0.50 \pm 0.01$ | $2.32 \pm 0.02$ | $0.53 \pm 0.02$ | $2.42 \pm 0.05$ | $0.41 \pm 0.02$ | $2.31 \pm 0.03$ |
| $m$ (kJ.mol<sup>-1</sup>.M<sup>-1</sup>) | $16.00 \pm 0.88$ | $6.65 \pm 0.39$ | $21.62 \pm 3.73$ | $7.48 \pm 1.13$ | $10.45 \pm 1.28$ | $6.55 \pm 0.42$ |
| $\Delta G^0_{H_2O}$ (kJ.mol<sup>-1</sup>) | $8.00 \pm 0.46$ | $15.43 \pm 0.93$ | $11.46 \pm 2.03$ | $18.10 \pm 2.76$ | $6.06 \pm 0.27$ | $15.13 \pm 0.98$ |
| $T_m$ (°C) | $46.5 \pm 1.7$ | | $50.9 \pm 0.7$ | | $44 \pm 0.1$ | |
| $K_m^{app}$ (mM) | $0.76 \pm 0.07$ | | $6.25 \pm 0.35$ | | $5.22 \pm 0.74$ | |
| $K_{cat}$ (sec<sup>-1</sup>) | $192 \pm 3.3$ | | $12.7 \pm 0.5$ | | $18.3 \pm 0.2$ | |
| $K_{cat}/K_m^{app}$ (sec<sup>-1</sup>.M<sup>-1</sup>) | $252.63 \pm 17.06$ | | $2.0 \pm 0.1$ | | $3.51 \pm 0.49$ | |

Enzyme kinetic analyses of the three PGM isoforms are also presented in Table 3. The catalytic efficiency of the PGM78, evaluated by the ratio *k*_cat_/K^app^_m_, appeared 125-and 70-fold higher than that of PGM90 and PGM100, respectively. Both changes in *K_m_^app^* (for the substrate Glucose 1 phosphate (G1P)) and *k*_cat_, explain most of the difference in the catalytic efficiency of the three isoforms. *K_m_^app^* (G1P) and *k*_cat_ of the PGM78 are indeed respectively tenfold lower and a tenfold higher than that of the two other isoforms (see Table 3).

### Fitness cost of individuals carrying the thermostable allele in terms of female fecundity

The Pompeii worm females exhibited an average coelomic fecundity of 200,000 oocytes with a great variability among them (values ranged from 1200 to 450 000 oocytes depending on size (age) and the reproductive state^[41]^). As opposed to our expectations, females carrying the allele 90 were on average more fecund than homozygous females carrying alleles 78 and 100. However, distributions of fecundity corrected by the size of the female were not significantly different one to each other according to *Pgm-1* genotypes (One-way ANOVA: F=1.08, p=0.37), see Fig. S5). This finding clearly indicates that the ability to live under hotter conditions is not counter-balanced by a lesser reproductive success, at least for the females.

## DISCUSSION

Based on allozyme data, Piccino *et al.*^[12]^ previously proposed that the enzyme polymorphism of the Pompeii worm *Alvinella pompejana* may be balanced at the locus *Pgm-1*, at least in populations of the northern EPR. They indeed showed that allozymes 90 and 100 display distinct thermal stabilities and kinetic optima, the frequency of the most thermostable isoform (allozyme 90) being positively correlated with temperature in newly-formed edifices^[12,36]^. Because the Pompeii worm is the only vent species able to colonize newly-formed still ‘hot’ hydrothermal chimneys, bearing thermostable alleles is likely to represent an adaptive advantage. The maintenance of polymorphism by selection on thermostable alleles is however a matter of questioning. If advantageous in the hottest part of the vent environment, thermostable alleles are indeed expected to rapidly spread in the population via recurrent selective sweeps. This is obviously not the case for the *Pgm-1* locus, which exhibited three major isoforms of different thermal stabilities (allozymes 78, 90 and 100) that sharply segregate across a barrier to gene flow depicted by Plouviez *et al.*^[44]^ at the Equator. Several hypotheses have thus been proposed by Piccino *et al.*^[12]^ about the maintenance of alleles at the PGM-1 enzyme. This includes (i) allele overdominance due to the rapid alternation of aerobic/anaerobic vent conditions, (ii) a fitness cost for individuals carrying the most thermostable allele at this locus, or (iii) a two-niches model effect due to fluctuating proportions of ‘hot’ and ‘cold’ habitats along the EPR.

In the present study, we sequenced the three major *Pgm-1* alleles of the worm to investigate the distribution of non-synonymous polymorphisms along the gene, examine their relationship with allozymes, and assess their evolutionary fate. We demonstrate that only two linked mutations (E_155_Q and E_190_Q) are associated with the net charge of allozymes and also responsible for the thermal performance of the three allozymes (78, 90 and 100). Looking at the evolutionary history of these alleles indicates they are part of a rather ‘old’ polymorphism that predates the vicariant event, which separated the EPR vent fauna across the Equator about 1.5 Mya^[44, 65, 66]^. In the following discussion, we therefore, examine arguments towards the adaptive maintenance of the PGM-1 isoforms, and proposed that thermal compensation represents a powerful mechanism by which different enzymatic properties might be maintained under balancing selection - at least, during the exploration of the mutational landscape of the protein that will lead to the emergence of the ‘optimal’ isoform - to optimize metabolic fluxes as previously stated by Eanes^[67]^.

### Two-allele polymorphism at the Pgm locus: a long story of balancing selection?

The non-synonymous polymorphism associated with the 4 allelic lineages of the *AP Pgm-1* appears to be quite low (π_N_=0.0025 on average). Such result sharply contrasts with earlier studies on branch-point glycolytic enzymes that control the metabolic flux for transport, storage and breakdown of carbohydrates, for which numerous cryptic non-synonymous changes have been described^[67]^. High levels of gene diversity were indeed observed between the slow, medium and fast electrophoretic *Pgm-1* alleles of *Drosophila melanogaster*^[14,68]^, or between phosphoglucose isomerase (PGI) alleles of *Colias* butterflies^[69]^ suspected to evolve under balancing selection. By contrast, only eight non-synonymous mutations (E37Q, V40L, E155Q, E190Q, R343I, G358S, T366M and F502L), some of which are at a low frequency, have been detected from the direct comparison of allelic sequences of *A. pompejana*. Moreover, even if gene diversity was slightly higher in the intronic region preceding exon3 and in the exonic region where the two EQ sites are found, its variation along the gene does not fit perfectly with the expectations of long-term balancing selection. A weak ‘hot spot’ signal of silent site variation supported by slightly positive Tajima’D and Fu&Li’F statistics is however observed near the doubly selected sites E^155^Q/E^190^Q but not as strong as signals depicted for the *Adh* locus in *Drosophila*^[70,71]^. Theoretical effects of balancing selection on nearby genome regions indeed promotes the increase of genetic diversity near the selected site due to the lack of recombination and the long-term accumulation of mutations^[38]^. The very low level of nucleotidic polymorphism found at the AP-*Pgm-1* can be however partially explained by recurrent population bottlenecks due to the challenging environmental conditions that affect the whole vent fauna^[15]^. The joint action of abrupt demographic changes and habitat specialization should indeed promote enzyme monomorphism. Under such conditions, the level of polymorphism observed at the *Pgm-1*, although low, appears to be quite unusual when compared to most of the genes examined in *A. pompejana*. In alvinellid worms, and especially thermophilic species, proteins are indeed under strong purifying selection with overall d_N_/d_S_ transcriptome means very close to zero with values ranged between 0.02 and 0.05^[72]^. To this extent, it is worth noting that gene diversity at the *Pgm-1* locus appears to be locally two-to four-fold higher than that recorded over the genome (ddRAD overall π=0.0025^[73]^) and from other reported genes^[35]^.

Looking more specifically at the 4 allelic lineages of the *AP-Pgm1* nearby the EQ sites (intron 2) clearly indicates that they have accumulated a great number of synonymous substitutions since their separation with almost no recombination events (low values of *R_m_* and *Rho*, see the Phase 2.1.1 analysis). The two allelic lineages L1 and L2 present in the Southern population exhibit 1% divergence between them, with a strong linkage disequilibrium between the two variant sites E^155^Q^190^ (PGM90) and Q^155^E^190^ (PGM78) and the silent substitutions found in intron2. This suggests that these two allelic lineages evolved separately without recombination in the vicinity of the two non-synonymous sites for a long period of time. The two northern allelic lineages (L3 and L4) also diverged by 0.7% divergence and two diagnostic indels in intron2. These silent mutations and indels are also linked together, suggesting once more that these two lineages evolved separately but for a shorter period of time. However, diagnostic mutations are not completely linked to the variant sites E^155^E^190^ (PGM100) and E^155^Q^190^ (PGM90). Although L3 forms a single clade associated with E^155^Q^190^, L4 is a mixture of E^155^E^190^ and E^155^Q^190^ suggesting either that, at least one recombination event occurred between the two northern alleles or that E^155^E^190^ (PGM100) is a recently derived variant in the northern population. Finally, both divergences observed between alleles within each population are of the same amplitude as the divergence estimated between the southern and northern alleles (0.9%), which coincide with an overall significant F_st_ value of 0.588 between them. If we accept that the southern (L1 and L2) and northern (L3 and L4) allelic lineages become isolated after the appearance of the physical barrier to dispersal, about 1.2 Mya^[35,44]^, this clearly indicates that the co-occurrence of the 4 highly divergent *Pgm-1* alleles derives from an older polymorphism predating the vicariant event that separated the Northern and Southern vent fauna of the East Pacific Rise, with the possible emergence of the genotype E^155^E^190^ in the North. Such a scenario is likely confirmed by the distribution of alleles in the exon3 haplotype network and the signature of the linked silent polymorphic sites in introns 1 and 2, where northern alleles seem to derive from a southern allele bearing the Q^155^E^190^ mutation.

### Selective modalities for the maintenance of a balanced polymorphism

The long-term evolution of *Pgm-1* alleles without recombination, at least in the first part of the gene, and their frequency changes according to environmental conditions raises questions about the selective modalities acting on the co-occurrence of alleles when one of the two alleles is better adapted to high temperatures^[12]^. One of the first explanations to the adaptive maintenance of AP-PGM1 isoforms was to consider that the rapid alternation of oxic and anoxic conditions during venting should favor heterozygote excesses if the heterozygote’s fitness is close to that of the favored homozygote in one of the two habitats^[74]^. Pogson^[13]^ previously proposed that overdominance represents the most likely evolutionary mechanism at the origin of the maintenance of a balanced polymorphism at the *PGM-2* locus for the oyster *Crassostrea gigas*, (but also see^[75]^ for its link with individual’s growth rate). Here, we were not able to detect any overdominance at the *Pgm-1* locus in terms of heterozygote excesses in natural populations, and simulations of structured coalescent with asymmetrical migration and overdominance always led to very high within-deme positive Tajima’s D and unequal diversities between demes not observed here. This finding confirms the previous study^[12]^ with allozymes. It is however worth noting that such advantage can be easily masked by the temporal dynamics of the thermal habitat (chimneys refreshing with time) and the juxtaposition of chimneys of different ages. The worm is indeed exposed to a mosaic of fluctuating thermal habitats where temperature could spatially vary according to the age of the chimneys^[12]^.

Maintenance of allozymes with different thermal stabilities can be also explained by a two-niches model of local differentiation with habitat and drift^[76,77,78,79]^. Simulating coalescences with a 2-niches model with a theta value equal to the observed value indeed provided parameters estimates (Fst, overall and intra-deme diversities with Tajima’s D) much closer to values observed in the vicinity of the two EQ sites than the two other models of asymmetric gene flow with and without overdominance. Given the spatial and temporal dynamics of the hydrothermal discharge, changes in the frequency of *Pgm-1* alleles could be either due to local adaptation or an exacerbated genetic drift associated with the dynamics of colonization of the newly opened sites. Indeed, the dynamic nature of hydrothermal vents over longer time scales (years) led to a very patchy and transient habitat, scattered along the EPR with a complex heterogeneity of age-driven vent conditions. This can be seen as a multitude of distinct ecological niches for the same species. In this context, the proportion of newly formed ‘still hot’ chimneys and older colder ones greatly varies over time depending on the spreading rate of the rift and thus the frequency of tectonic and volcanic events along the East Pacific Rise.

Piccino *et al.*^[12]^ also proposed that the maintenance of a bi-allelic polymorphism at the *Pgm-1* locus results from a fitness cost to the colonization of the early stages of a chimney. The first settlers on ‘hot’ (>100°C) anhydrite chimneys indeed benefit from a lack of predators and competitors. Colonists may therefore display more thermoresistant alleles but a lesser reproductive investment and/or survival. Watt and collaborators^[80,81,82]^ and Wheat *et al.*^[69]^ previously showed that both PGI and PGM have a great contribution on the male mating fitness in *Colias* butterflies, probably as the result of longer and more vigorous flight within the day. Based on female fecundity, we were however not able to observe fitness differences between the *Pgm-1* genotypes. This suggests that a better predisposition to colonize still ‘hot’ chimneys is probably not compensated by a reduced reproductive success to prevent the fixation of the advantageous allele. In the fruitfly, the *Pgm* locus represents however a quantitative trait for glycogen storage and hence, the ability to survive better to starvation^[34]^. In the case of *A. pompejana*, differences in the thermal regime could be great between colonists and reproducers. As a consequence, colonists subjected to longer periods of high temperature (and associated hypoxia) may be maladapted to produce and use glycogen reserves and, thus, may not invest much into reproduction. On the contrary, secondary settlers arriving in much cooler conditions are more likely to use their glycogen reserves to massively invest in the production of gametes as previously shown by Faure *et al.*^[41]^. Colder conditions indeed seem to be a prerequisite for releasing fertilized eggs after pairing as embryos are not able to develop at temperatures greater than 15°C^[40]^.

### Adaptive polymorphism: a trade-off between enzyme thermostability and catalysis

In *Drosophila*, *Pgm-1* variants play a non-negligible role in the regulation of the metabolic energy pool along latitudinal clines where the decrease of temperature is compensated by an increase of the enzyme activity. Populations of *Drosophila melanogaster* living at the highest (and thus coldest) latitudes possess PGM allozymes with a higher catalytic efficiency and greater glycogen contents. According to the theory of metabolic flux, this can be an adaptive way of temperature compensation to maintain the same glycogen contents over the latitudinal gradient^[67]^. Differences in both protein thermostability and catalytic efficiency between *Pgm* alleles were previously reported to explain both local differentiation and latitudinal clines in the oyster *Crassostrea gigas*^[6,13]^ and *Drosophila melanogaster*^[34]^. By comparison, thermal compensation may be therefore directly linked to different biochemical phenotypes that interact with the growth rate and reproductive effort of the worms. The theory predicts that differences in activity at only one enzyme must be however substantial to affect metabolic fluxes between genotypes^[83]^. To test this hypothesis, one could measure the effect of non-synonymous mutations on the functional properties of the enzyme, its conformational stability and their effect on population fitness. In this study, three recombinant isoforms of PGM-1 (E^155^E^190^ (PGM100), E^155^Q^190^ (PGM90) and Q^155^E^190^ (PGM78)) were obtained by directed mutagenesis. The replacement of the glutamate by a glutamine at position 190 increases the conformational stability and thermostability of the protein, confirming that PGM90 is the most thermostable isoform. As discussed by Piccino *et al.*^[12]^, carrying this allele may be advantageous during the colonization of newly-formed chimneys whose surface temperature usually exceeds 50°C. Unexpectedly, PGM90 exhibits a decrease in the catalytic efficiency of the enzyme when compared to the two other recombined variants, (k_cat_/K_m_ is a hundred times for PGM78 and nearly two-fold greater for PGM100 when compared to PGM90). The recombinant PGM90 also exhibits the lowest affinity for its substrate glucose-1-phosphate at 17°C. This finding is of importance because such a genetically-determined trade-off between protein stability and enzyme activity was not reported in other invertebrate species subjected to balancing selection so far^[6,13,34,82]^. Increased thermostability of a protein is often associated with a decrease in the flexibility of the molecule, and thus the dynamics of the enzyme reaction^[84,85]^. Our results are in perfect agreement with these theoretical expectations and support the positive role of a thermodynamic trade-off between thermostability and catalysis as previously proposed by Eanes^[11]^ to explain the co-occurrence of alleles. PGM90 can remain stable for a longer period of time but is less efficient to either produce or consume the glycogen reserves of the worm than the two other isoforms are. To this extent, the fact that the k_cat_/K_m_ ratio of isoform PGM78 is much higher than that of the isoform PGM100 can explain why isoform 78 is more frequent in the Southern populations (about 80%) when compared with isoform PGM100 in the Northern populations (around 70%). The balance between allele frequencies from both sides of the Equator may be dictated by the selective coefficient attributed to each genotype as the direct reflection of the catalytic efficiency difference between PGM90 and its alternative isoforms.

### Structural effect of mutations E155Q and E190Q

The location of the two main polymorphic sites (E155Q and E190Q) onto the 3D model structure of the phosphoglucomutase 1 are exposed to solvent and not located in the binding domains of the enzyme (Fig. 6). Their potential effect on the catalytic properties of the enzyme is therefore not the result of a direct interaction with the substrate and/or the residues involved in the catalysis. This is not surprising as most of the mutations affecting the binding of the substrate glucose-1-phosphate, the Mg^2+^ ion and the phosphate should be deleterious. Similarly, in a study of polymorphism of the *PGM* from *D. melanogaster*, none of the 21 polymorphic amino-acid replacements were located in the catalytic site of the enzyme^[34]^. Based on their location, both substitutions should affect the net charge of the protein in the same way. However, the isoforms 78 and 90 do not have the same electrophoretic mobility, suggesting that some post-translational modification may be involved in the electrophoretic separation of the three isoforms. The gain of a glutamate at position 155 may be associated with a potential ionic bond with histidine 157 with a distance of 6.5 Å between them. Ionic and hydrogen bonds have been shown to increase the stability of enzymes^[86]^, and partially explain the thermostable 3D structure of the Cu-Zn and Mn superoxide dismutase enzymes in *A. pompejana*^[87,88]^. This may account for the increased thermostability of isoform 90 but should also have the same effect for isoform 100, which is obviously not the case. This suggests a negative effect of glutamate (E) at position 190 that would negate the positive effect of glutamate at position 155. Alternatively, the 3D model comparison of the three isoforms shows that the replacement of one glutamine (Q) by a glutamate (E) at position 190 (allozymes 78 and 100) introduces a negative charge in a region already enriched in acidic residues. This high density of negative charges could have a destabilizing effect on the protein structure by a Coulomb repulsion effect and could thus lead to greater sensitivity to temperature. Finally, the glutamine replacement at position 155 is also likely to play a key role in the molecular dynamics of the protein, especially during the 180° rotation of the reaction intermediate (glucose-1,6-diphosphate) inside the active site. This likely explains the higher enzymatic efficiency of the isoform 78.

**Fig. 6.**
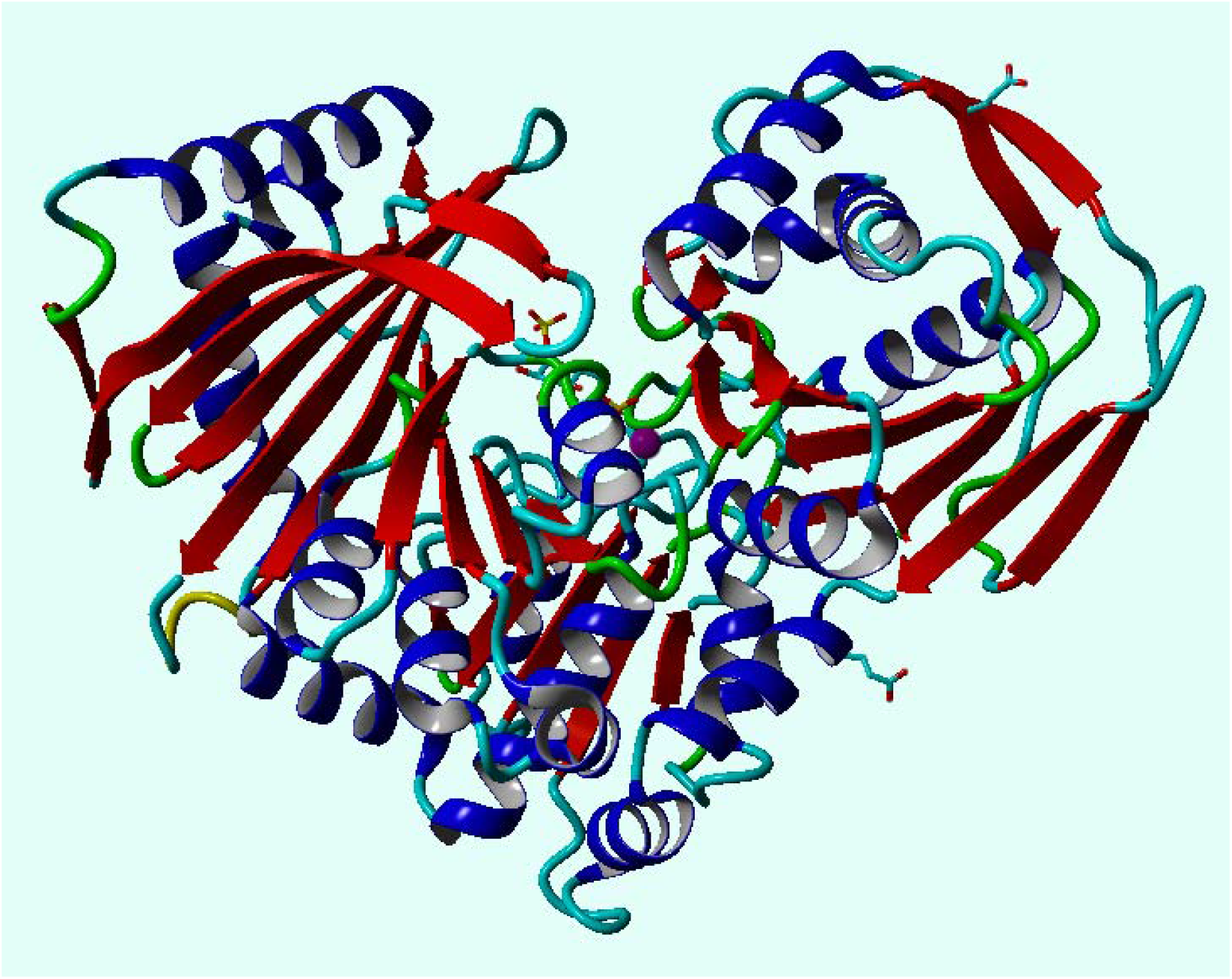
3D structural model of *A. pompejana* PGM 78 fitted on the PGM-1 rabbit template (1C47, 2.70Å) using Modeller 9v13. The protein is structured in 4 domains labelled from I to IV (I green, II yellow, III blue, IV violet). Positions 155 and190 of EQ replacements belong to domain I near to catalytic site of the enzyme, which binds the reaction catalyser, alpha-D-glucose-1,6-diphosphate, and the ion Mg^2+^.

## Acknowledgments

This article has been written in memory of Dominique Le Guen, who greatly helped us in setting up of the allozyme genotyping. We thank chief scientists and the ‘Nautile’ crews for their technical support and efforts during the oceanographic expeditions Phare2002, Biospeedo2004 and Mescal2010. We are very grateful to the two anonymous referees who provided valuable comments and editorial suggestions on the manuscript.

