## Supplemental Tables and Figures for "Balanced polymorphism at the *Pgm-1* locus of the Pompeii worm *Alvinella pompejana* and its variant adaptability is only governed by two QE mutations at linked sites"

Table S1. Sequence of the primer pairs, and their use in the study. Primer names are based on the human *Pgm-1* exon number, but do not always correspond to the exon/intron nomenclature in *A. pompejana*, due to the comparative fusion of some exons (case of AP-exon3 with corresponds to exon2, exon3 and exon4 in human). Lower-case sequences correspond to intronic regions and upper-case sequences correspond coding regions.

| **Primers** | **5' Sequence 3'** | **Localization/Method** |
| --- | --- | --- |
| Anchor-OligodT | 5-CTCCTCTCCTCTCCTC-T(17)-3 | Primer for reverse-transcription of polyA+ mRNA |
| AP_PGMcDNAF1  AP_PGMcDNAF2  AP_PGMcDNAR | 5-GTNGTYGGNGGNGAYGGNBG-3  5-YCAYAAYCCNGGNGGNCC-3  5-NGTRATNACNGTNGGWKC-3 | Pgm-1 cDNA fragment amplification |
| Ap_PGMex1F  Ap_PGMex2R | 5-AAG AGG CAT CAG AGA AGA TA-3  5-GAC CAC CTG GGT TAT GAG AT-3 | Gene structure determination (amplification of exon1-intron1) |
| Ap_PGMex2F  Ap_PGMex4R | 5-TAG GTA AAG ATG GCA TAC TT-3  5-CCG CTG AGA GCA TTG ACA AG-3 | Gene structure determination (amplification of exon2-exon3) |
| Ap_PGMex5F  Ap_PGMex7R | 5-AA GAC TTT GGA GGA GGA CAT C-3  5-ATC CAT AAG GTT ACC AAA GAA CTT CCA-3 | Chromosome walking along the gene (exon3-exon5) |
| Ap_PGMex7F  Ap_PGMex9R | 5-AGT TCT TTG GTA ACC TTA TGG ATG CT-3  5-ATT TTA TCA AGG TTG GCC ATC ATC TG-3 | Gene structure determination (amplification of exon5-intron6) |
| Ap_PGMex9F  Ap_PGMex10R | 5-AC CTT GAT AAA ATG GCA GCT GAC-3  5-GA ATC TGA CTC ATA GCT ATC AAT GTA-3 | Gene structure determination (amplification of exon7-exon8) |
| Ap_PGMex10F  Ap_PGMex11R | 5-ATG TAC ATT GAT AGC TAT GAG TCA GAT-3  5-AAA TTT AAG TAA TAA CAG TAG GCT GCT-3* | Gene structure determination (amplification of exon8-exon9)  **anchored with the stop codon* |
| Ap_PGMint1R1  Ap_PGMint1R2  Ap_PGMex1Rn | 5-tgtgtatataagacgttctttttactgtgg-3  5-tcatgtaaaattcttgatatacttacacc-3  5-GTA GCC TTT CTC AGA CCA CTA GT-3 | Chromosome walking to obtain the 5’UTR and the first exon with w1, w2, w3 and wc (Mishra et al., 2002) |
| Ap_PGMint4F  AP_PGMex5R | 5-tgttaggtagcatgcctcca-3  5-TCC TCC AAA GTC TTC CAG TG-3 | Genotyping EQ mutations in exon3 |
| Ap_PGMex1F  Ap_PGMex2R | 5-AAG AGG CAT CAG AGA AGA TA-3  5-GAC CAC CTG GGT TAT GAG AT-3 | Genotyping intron1-exon2 |
| Ap_PGMex6F  Ap_PGMint7R | 5-ttt tag GAC AGA AAC ATG ATA CTT GGT-3  5-taatattac CT GAT GTG ATC TGA TCC-3 | Genotyping intron4-exon5 |
| Ap_PGMex9F  Ap_PGMex10R | 5-AC CTT GAT AAA ATG GCA GCT GAC-3  5-GA ATC TGA CTC ATA GCT ATC AAT GTA-3 | Genotyping exons 6 and 8 |
| AP_PGMmut78F  AP_PGMmut78R | 5-AAG TGA TAG ACT CTG TGC AGG ATT ATA TGG AC-3  5-TAG TCC ATA TAA TCC TGC ACA GAG TCT ATC AC-3 | Directed mutagenesis to produce allele 78 from allele 100 |
| AP_PGMmut90F  AP_PGMmut90R | 5-TTG ACA CAA AAG ATC ACA CAA TAC CAC ACT GTG C-3  5-AGG CAC AGT GTG GTA TTG TGT GAT CTT TTG TGT C-3 | Directed mutagenesis to produce allele 90 from allele 100 |
| Pet20_PGMXhoI  Pet20_PGMAseI | 5-CTC GAG AGT AAT AAC AGT AGG CTG CTG TC-3  5-ATT AAT GAG TCT GAA GTC GGT GAC AGT GGC T-3 | Cloning in Pet20 overexpression vector |
| PetDuet_PGMBamHI  PetDuet_PGMNotI | 5-GGA TCC GAG TCT GAA GTC GGT GAC AGT GGC T-3  5-GCG GCC GCT TAA GTA ATA ACA GTA GGC TGC TGT C-3 | Cloning in PetDuet overexpression vector |

Table S2. Frequencies of EE, EQ and QE *Pgm-1* alleles, heterozygosities and Fis (*: significant with 1000 permutations) in northern and southern populations of *Alvinella pompejana*. Note: Allele QQ was not found in any of the populations.

| **Site** | **N** | **EE** | **EQ** | **QE** | **Hobs** | **He_(n.b.)_** | **Fis** |
| --- | --- | --- | --- | --- | --- | --- | --- |
| South EPR overall | 126 | 0.051 | 0.139 | 0.810 | 0.293 | 0.324 | +0.094 |
| Krasnov (21°33’S, hot) | 23 | 0.044 | 0.217 | 0.739 | 0.434 | 0.413 | -0.053 |
| Bordreaux (21°25’S, hot) | 30 | 0.033 | 0.100 | 0.867 | 0.267 | 0.242 | -0.105 |
| Fromveur (18°25’S, hot) | 32 | 0.000 | 0.047 | 0.953 | 0.094 | 0.091 | -0.033 |
| Rehu Marka (17°25’S, cold) | 41 | 0.110 | 0.195 | 0.695 | 0.390 | 0.472 | +0.176 |
| North EPR overall | 94 | 0.718 | 0.277 | 0.005 | 0.404 | 0.410 | +0.014 |
| Jumeaux (13°N, hot) | 19 | 0.605 | 0.395 | 0.000 | 0.684 | 0.491 | -0.410* |
| Julie (13°N, cold) | 28 | 0.696 | 0.286 | 0.018 | 0.429 | 0.441 | +0.028 |
| Genesis (13°N, cold) | 27 | 0.741 | 0.259 | 0.000 | 0.296 | 0.391 | +0.246 |
| Elsa (13°N, cold) | 20 | 0.825 | 0.175 | 0.000 | 0.250 | 0.296 | +0.159 |

Table S3. Summary statistics of structured coalescent simulations (n=1000) obtained with the msms solftware and the pylibseq librairies in order to test a model of asymmetric migration across a barrier with and without selection and two levels of recombination in order to examine the genetic expectations of overdominance and the two-niches (2 demes/2 habitats) models. *: selection coefficient was set to 100 (low=ls) and 10 000 (high=hs) for the selection models. For comparison purpose, the within deme values correspond to parameters estimated for the exporting deme in the migration asymmetry. Values between brackets represent the confidence interval of the parameter’s simulations at 95%. D: Tajima’s D, AM: Asymmetric migration across a barrier with no selection, AM_OD: Asymmetric migration across a barrier with overdominance, AM_2N: Asymmetric migration across a barrier with a two-niches model.

| Selected model | R | Overall π | Overall θ_W_ | Overall D | Fst | Within deme π | Within deme θ_W_ | Within deme D |
| --- | --- | --- | --- | --- | --- | --- | --- | --- |
| Obs. data* | **1** | **8.2** | **8.4** | **-0.12** | **0.27** | **8.0** | **7.6** | **+0.20** |
| AM | 0 | 34.2  [32.6, 35.9] | 17.1  [16.5, 17.7] | +2.73  +2.6, +2.8] | 0.85  [0.83, 0.86] | 3.1  [3.0, 3.2] | 3.1  [3.0, 3.1] | -0.05  [-0.1, +0.0] |
| AM | 1 | 34.4  [32.6, 36.3] | 17.1  [16.4, 17.8] | +2.65  [+2.6, +2.7] | 0.84  [0.83, 0.85] | 3.1  [3.0, 3.3] | 3.1  [3.0, 3.2] | -0.04  [-0.1, +0.0] |
| AM | 100 | 34.8  [33.1, 36.5] | 17.3  [16.7, 18.0] | +2.75  +2.6, +2.8] | 0.85  [0.84, 0.86] | 3.1  [3.0, 3.2] | 3.1  [3.0, 3.1] | -0.04  [-0.1, +0.0] |
| AM_OD-ls* | 1 | 37.0  [35.4, 38.6] | 19.5  [18.8, 20.1] | +2.59  [+2.5, +2.7] | 0.73  [0.72, 0.75] | 6.4  [6.2, 6.5] | 4.6  [4.5, 4.7] | +1.20  [+1.1, +1.3] |
| AM_OD-hs* | 1 | 37.0  [35.5, 38.6] | 19.4  [18.7, 20.0] | +2.61  [+2.5, +2.7] | 0.74  [0.73, 0.75] | 6.3  [6.1, 6.4] | 4.5  [4.4, 4.6] | +1.21  [+1.1, +1.3] |
| AM_OD-hs* | 100 | 35.1  [34.3, 35.8] | 17.6  [17.3, 17.9] | +3.20  [+3.1, +3.3] | 0.90  [0.89, 0.90] | 3.2  [3.1, 3.3] | 3.1  [3.0, 3.2] | +0.05  [-0.0, +0.1] |
| AM_2N | 1 | 21.8  [21.1, 22.6] | 16.9  [16.5, 17.3] | +0.83  [+0.7, +0.9] | 0.45  [0.44, 0.47] | 14.6  [13.8, 15.3] | 12.6  [12.2, 13.1] | +0.44  [+0.3, +0.5] |
| AM_2N | 100 | 22.7  [22.4, 23.0] | 17.4  [17.3, 17.6] | +0.98 [+0.9, +1.0] | 0.50  [0.49, 0.51] | 15.0  [14.7, 15.2] | 12.8  [12.6-13.0] | +0.58  [+0.5, +0.6] |

**Fig. S1.** Entire sequence of the *Pgm-1* gene of *A. pompejana* and its translated exons

(1) Structure of the *Pgm-1* gene*.* Grey zones represent positions where forward and reverse primers have been designed. Exons are indicated by the use of uppercase and introns are in lowercase. Highlighted codons in yellow* represent polymorphic non-synonymous changes between alleles found at a frequency of more than 10%, and in green below 5% but which are likely to change the net charge of the protein. ^#^ symbol represents methylated codons found at the end of exon 5.

ATGAGTCTGAAGTCGGTGACAGTGGCTACGAAGCCCTTCGATGGGCAGAAGCCGGGCACTAGTGGTCTGAGAAAGGCTACGAAGATATTTATGCAAGAACATTACACA**GAA**AACTTC**GTG**CAATGTACGTTGTCTGCCATGGGCGACAAATTAAAGGGATGTACACTAGTAGTTGGAGGTGATGGAAGGTATTATGGTAAAGAGGCATCAGAGAAGATAATTAAAATGTGCGCAGGTAATGGT**gt**aagtatatcaagaattttacatgatgtgtaaacaattgtttctaactgacatcagaagccacagtaaagagaacgtcttatatacacagataatatatgttaacacaactttcttctcgttgcattaaatgcggtgattaatttttgctttatttgaaactaccaagtttatttaacgtatttatttgttcacaatttgaaatatgtttatactgtttacatgcctgaatttggtttgaattgacgtttttaaatgtactaattacgttttgttgttgttgtttttttagactatttgaatcatttaaccagccatgctaattttgattctattgtttgtgcttattacatttgtaccagatatgaaaggggagttaagaatgtgttgtaccatgatttgtatggtgatttaacattaatgcaattatgtttgtgtatttatatat**ag**GTAGCAAAGGTAATTATAGGTAAAGATGGCATACTTTCTACACCAGCTGTGTCATGCTTGATCAGAAAAAATCACACTGATGGAGGAATAATCCTCACTGCATCTCATAACCCAGGTGGTCCAAATGCTGATTTTGGCATAAAGTTTAATATTGCCAACGGAG**gt**aattcattcatgtttctacctgttcaaatcctttaaaccatacaaacaaatccttctatttgcaagaccaataaaaatgttcaatgtttaatcaatggtaacagctgaatgtaagtgtatgtatttttcaaatgtgatcaatataatataaatggactattttaaagacagtagagtctttctatgattgccatgtttttaattgattctgtctttttaaccatctgttgtattaaggggtgaaactagtattctgcataatgctatatgtttattcattatctgttagatttgaagagaataatatcataatagttagttaacctttttatttttagtagtgtttgtttcattaaaatggcacatcagctcattttgtagtgatttttatcatttgtgtagtttatatttattaaagataaataatactctctacttttattagttgtagttgttttgggggtctatacaccaaactgttcatatcatgataactttatgactatgggtgttatgtcaccatgatagaaacttgatatttaaagagcaaagtgaactagtagatccagattccacagttggtcctggttaaaatattcaacactgtctgtcactgatgtacacaaattgttgtgatccatggatgttctataaatgtttgcttagcatttgatgtaatatatgctacatagtttttgctgtatctactgcctgtccttttccatattttattgggactacttggtcactttagctgaccaaggacttgttaggtagcatgcctccattatttgtatgattaatgtaggtttgaaagctgaccaatgttaatatttttcatcac**ag**GACCAGCCCCAGCTGGAGTCACAGATCACATCTATGCCTTGACACAAAAGATCACA**GAA**TACCACACTGTGCCTGACCTGAAGGCTGACATCTGTACAATAGGAAGCCAGACGTTTACTGTTGATGATCATCCATTTAATATTGAAGTGATAGACTCTGTG**GAG**GATTATATGGACTACATGAAGGAAATTTTTGACTTCAATTCCATCAGAGGCTTATTGACTGGAGAAGGAGGACAAACAAAGCTAAAGGTCCTTGTCAATGCTCTCAGCGGAGTGGTTGGCCCATATGTGAAGAGGATACTGTGCCAGGAGCTAGGCATGGATGAAGCCAGTGCTGTTAATTGTGTTCCACTGGAAGACTTTGGAGGAGGACATCCAGATCCCAACTTGACCTATGCAGCTGATTTAGTGAATGAATTAAAGAAAGGTGTCCATGATTTTGGTGCTGCATTTGATGGC^#^GAC^#^GGC^#^**gt**aagttaattagtttgtgatagatgtcttattatttttatggtaaaactgacaagtatagctagaaaaatactttcaaaggatgtatagttctagaattgttgataaaaaacaattcattatttcagttgcttatatatttgtttgtgtggctgaggtgctgattgtctcacatctcaaaatgttttggtgtttggtgactttt**ag**GACAGAAACATGATACTTGGTAAGAATGGCTTCTTTGTATCGCCATGTGACTCCCTGGCAGTCATAGCTGCACATTTGGAGTGTATACCATATTTCAAGAAGTCTGGCATAAAAGGTTATGCC**AGA**AGCATGCCAACTAGTGGGGCCATTGAT**AGgt**aaatatataaaaatgatcatctgtggacctaattctgttatgtttttcagttcttttgttttaccttacttcatctcatatatagtagtcacagatgatatattgatcatcttattctgaactaggctacaaaataagagtttgccagtctaataaattaaaattatcacactaactgctgctttttatgatac**agA**GTGGCACAGAAGAAGGGCAAGGAGATGTTTGAGGTGCCAACAGGCTGGAAGTTCTTTGGTAACCTTACGGATGCTGGTAGACTTTCACTTTGTGGAGAGGAGAGCTTTGGGACAGGATCAGATCACATCAG**gt**aatattacacaatgctacatgtgtcaaatttcattagaaataacttcatcttgtaagttgtatacactttctgtaagccgactagagtttccatttcagaattcatttaacatgttatcagtcatatatattctatatacggatttgaattttgtaatgtgagctatccatagctgatattgacatattgctgctcagttgcctatatgagactaaggcagtcttatctctcattcttggatgctgctatctataaaaccattttgacataatacatcatcaaagtgtgtgtttgtattacatcattatgtgttgaaatgttgaactgtttattgtgtttc**ag**A**GAA**AAGGATGGCTTATGGGCAGTTTTGGCCTGGCTCTCTATTTTAGCCTGCAAGAAGCAGTCAGTAGAAGAAATCTTAAAGGATCACTGGAAGACATATGGCAGGAACTTCTTCACTAG**gt**taatatcatagatgtattacagtctcataaaatcatgtataaaatttataatatttaatcacagttttgctgtaataatatgaaaacctacttgaaattgataagatgtgaaaaataactttgtggctgttacttgtaccttctttatttgttgtggtgactgacagttgcacctatttgc**ag**GTATGATTATGAGAATGTTGAATCTGATCCAGCCAATCAGATGATGGCCAACCTTGATAAAATGGCAGCTGACTCATCTATTGTTGGCAAGGTGTTCAGTCATGGTGACAAATCATATAAAGTAGCCAAGATGGACAACTTTGAATACACTGACCCAATTGACAACAGTGTATCAAAAAAACAG**gt**aattctgtcatcttattaattgtgcacacatacacacctttgttcacatttgtgttaattatttgctctgtaaaataaatactgcccaattcagaatgaaaaaaaaatgtctttcttacatcatgcaattgttattttgtgtcacctatat**ag**GGCATCCGGATCATTTTTGAGGATGGATCAAGGATTATATTCCGTCTGAGTGGTACAGGAAGTGCTGGAGCAACAATCAGGATGTACATTGATAGCTATGAGTCAGATTCAAACAAACAGCTCCTAGATTCTCAG**gt**ttgttcacagcttaaatataacaagtgttatatattctaaatttgcagcattgatgctcaatattgtgattaatattttagatataattttatattatgaaagaccttatcattaccactgcttatgaactgattgtttatgtattaaagtagtttcttgagtggttattagggaaggcatgatgtcacttcattttcattctaatgttagcactatataaaatgctttaaaaatctatgcacaactcccactgaacaaatatttatttaattatcattccattctgc**ag**GTCATGCTGAAACCACTGATTGAGATAGCACTGGAAATATCCCAGCTTAGAGAGCTGACAGGAAGACAGCAGCCTACTGTTATTACTTAAatttggtatgcagccaacatttttgtcttcaattaccatgtagtgctatgtcatgtgatgagctatgcttagatgatctgtatgtaaaaaaaaaaaaaaaaaaaaaaaaaaaaaaaaa

(2) translated cDNA sequence of the AP-PGM enzyme (562 aa). Bold uppercase letters between parentheses are the three polymorphic sites for which the alternative allele has a frequency greater than 5%. Bold lowercase letters correspond to alternative mutations, which frequency is lower than 5% in genotyped regions. Other bold sites represent non-synonymous singletons in our set of sequenced individuals and, grey zones correspond to intronic regions linking the translated regions. *: stop codon.

MSLKSV**T**VATK**P**FD**G**QKPGTSGLRKATKIF**M**Q**E**HYT**(E/q)**NF**(V/L)**QCTLSA**M**G**D**KLKGCT**L**VVGG**D**GRYYGKEA**S**EKIIKMC**A**G**N**GVAKVIIGK**D**GILSTPAVSCLIR**K**NHTDGGIILTASHNPGGPNADFGI**K**FNIANGGPAPAGVTDHIYALTQKI**T(E/Q)YH**TVPDLKADICT**I**GSQTFTVDDHPFNIEV**ID**S**V(E/Q)D**Y**MD**YMKEIFDFNSIRGLLTGEGGQTKLKV**L**VNALSGVVGPYVKRILCQELGMDEASAVNCVPLDFGGGHPDPNL**TY**AADLVNELKK**G**VH**D**FGAAFDGDGDR**(N/d)MI**LGKNGF**F**VSPCDSLAV**I**AAHLECIPYFKKSGIKGYA**R**SMPTSGAID(**R/i)**VA**Q**KKGKE**M**FEVPT**(G/s,d)**WKFFGNL(**M/t)**DA**G**RLS**L**CGEESFGTGS**D**HIR(**E/k)**KDGLWAVLAWLSIL**A**CKKQSVE**E**ILKDHWKTY**G**RNFF**T**RYDYENVESDPANQMMANLDK**M**AADSSIVGKVFSHGDKSYKVAKMDNFEYTDPIDNSVSKKQGIR**(I/l)**IFEDGSRI(**I/t)(F/l)**RLSGTGSAGATIRMYIDSYESDSNKQ**L**LDSQV**M**LKPLIEIALEISQLRELTGRQQPTVIT*

**FigS2.** Haplotypes network obtained by genotyping of the *Pgm-1* exon 3 on 187 individuals coming from the North and the South EPR. The red and the yellow correspond to the populations of the North and the South, respectively. The numeric values are the numbers of sequences forming each haplotype. Mutations non-synonymous and the concerning amino acids are presents on the branches.

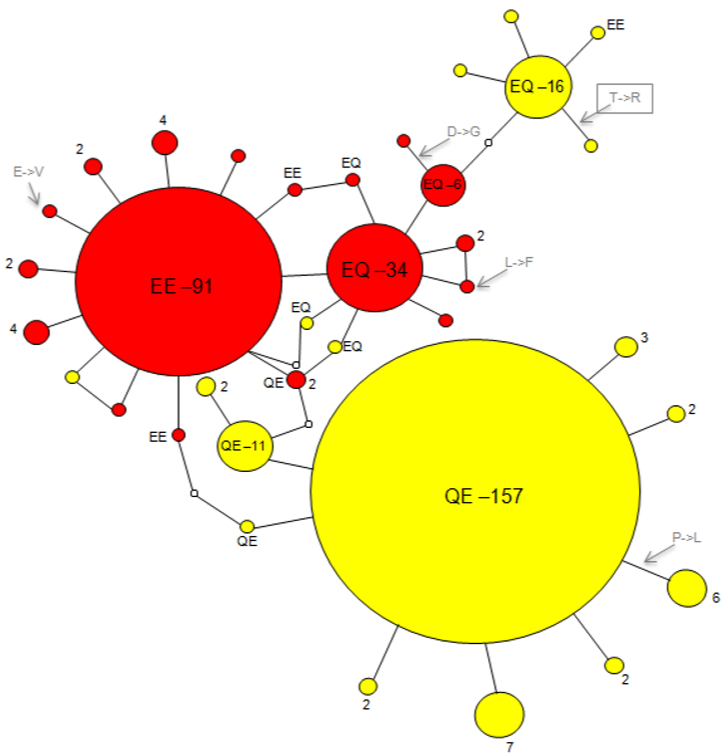

**FigS3.** Guanidium chloride (GdmCl) denaturation curves for the three PGM1 overexpressed isoforms. The fluorescence intensity at 324 nm (excitation at 290 nm) is presented as a function of GdmCl concentration. (A) Evolution of the protein denaturation along the GdmCl gradient of isoform QE (78) showing the normal (N), intermediate (I) and denaturated (D) states of the protein. (B) Denaturation curves *f_u_ (I)* and *f_u_ (II)* of the three isoforms showing that isoform differences stand during the first step of protein denaturation.

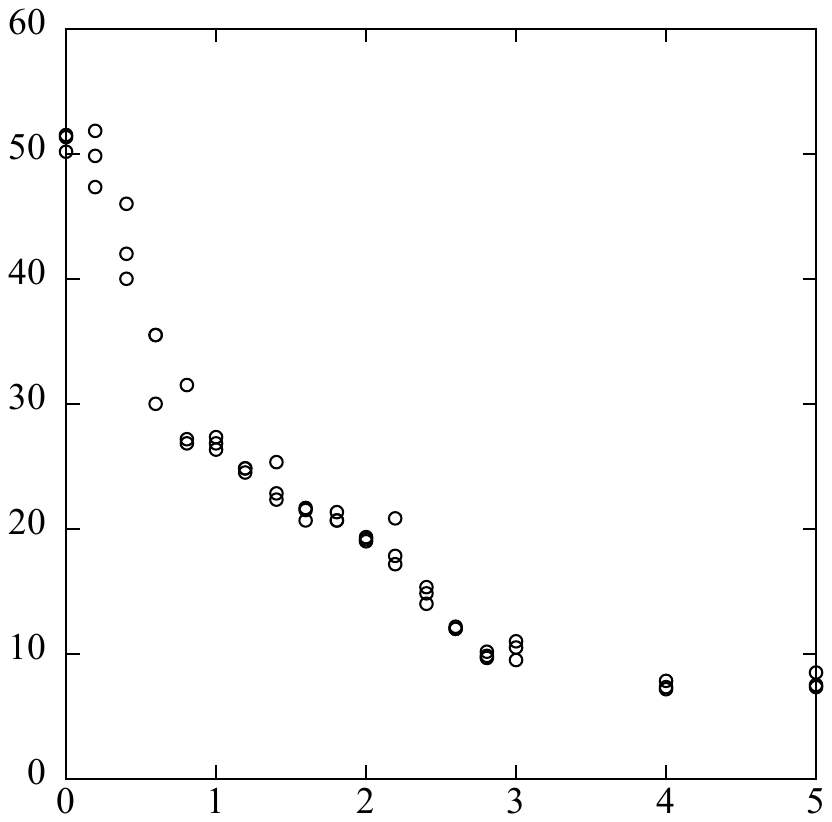

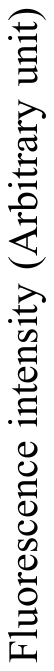

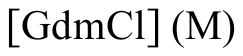

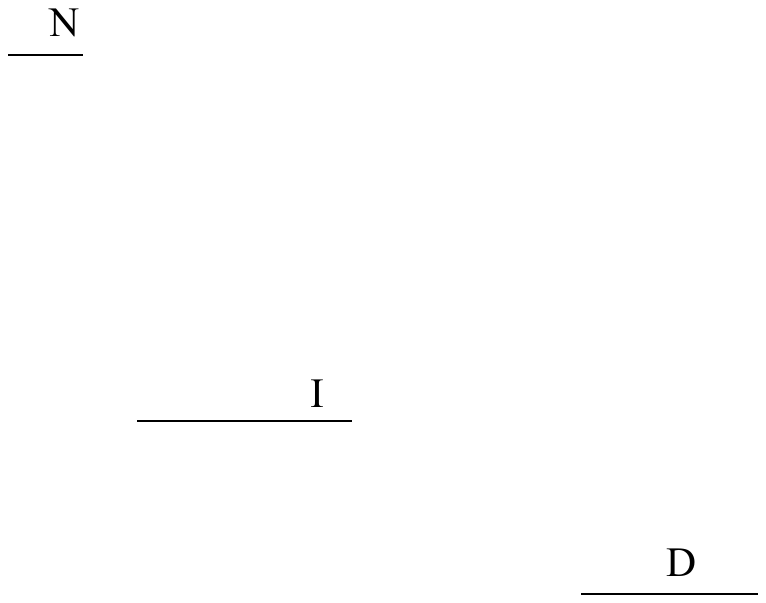

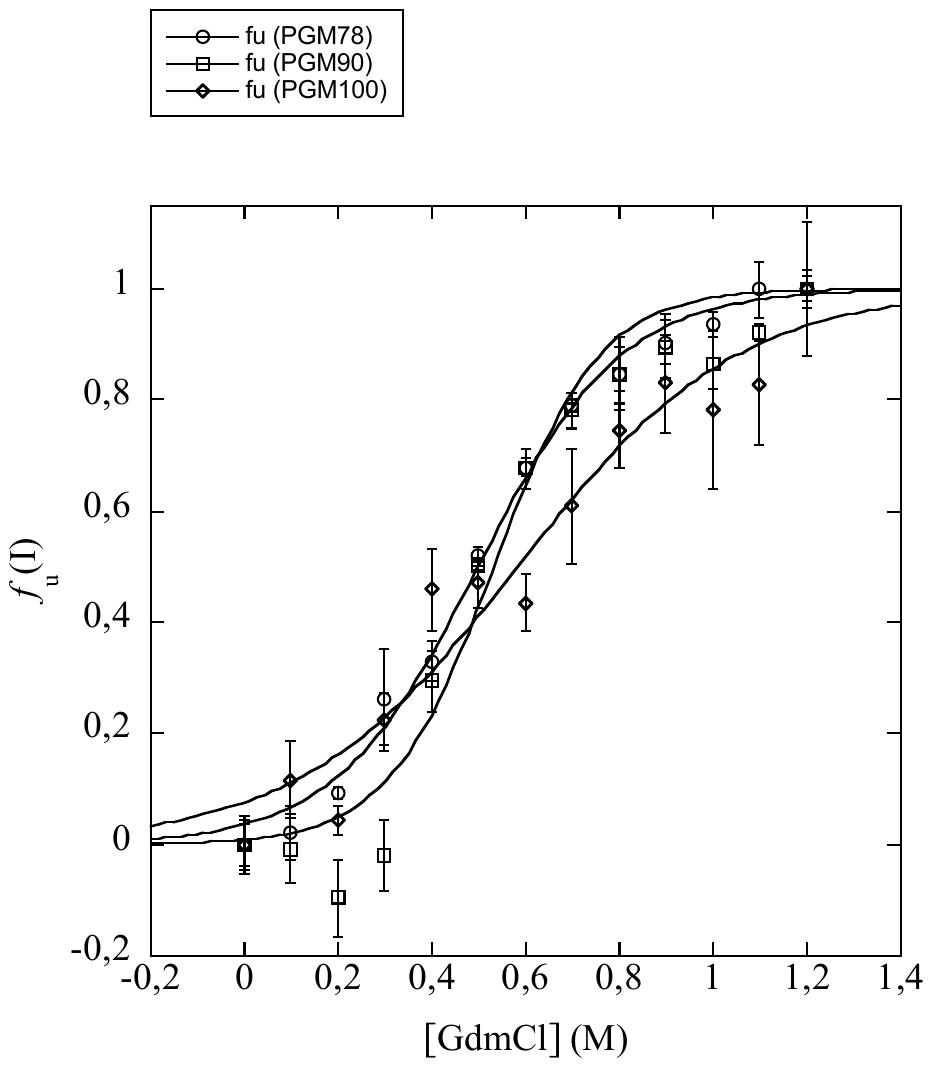

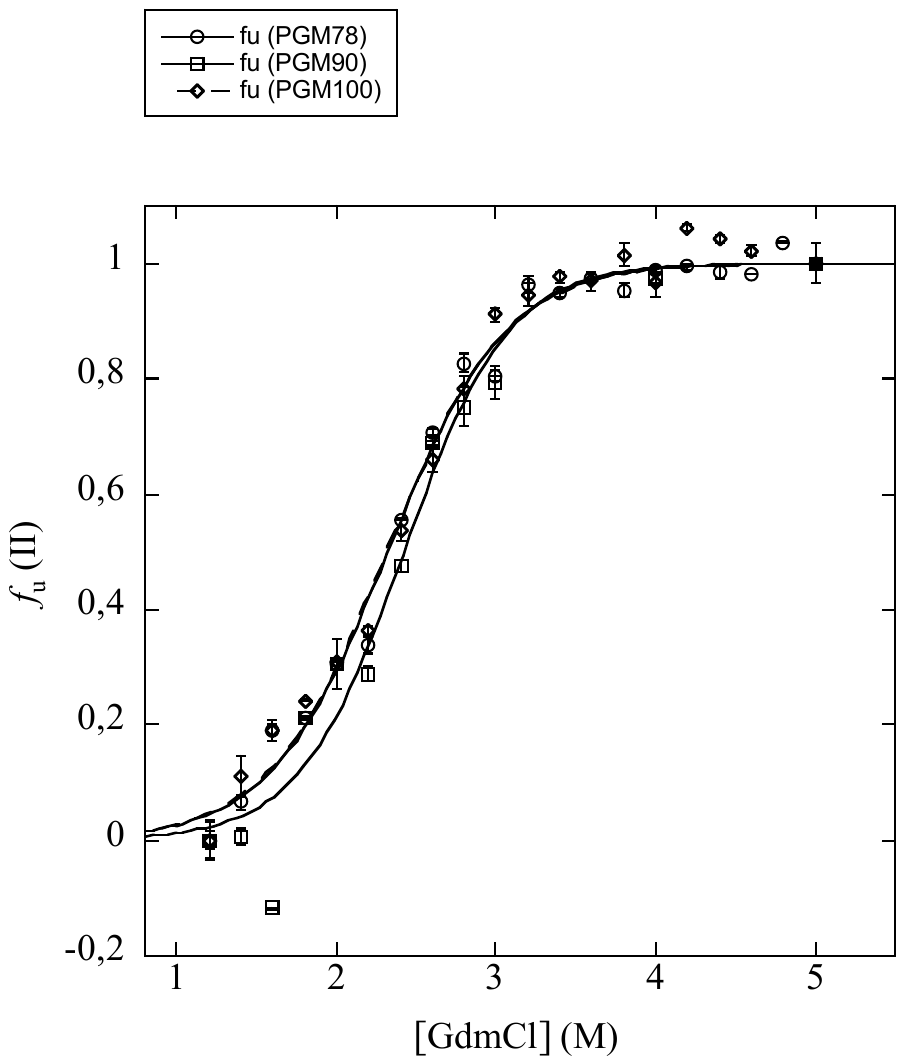

**Fig S4.** Modelled regression curves of the folded/unfolded protein states *f_u_*(I) and *f_u_* (II) of three PGM1 overexpressed isoforms for each transition according to the denaturation equilibrium N↔I↔U (Native↔Intermediate↔Unfolded).

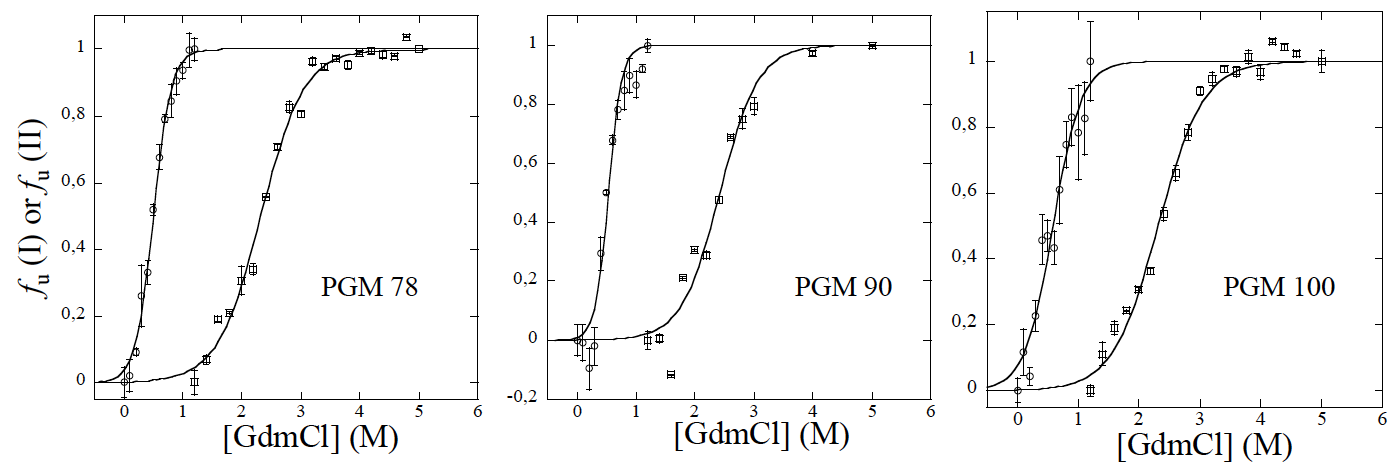

**Fig. S5.** Scatterplots of female *A. pompejana* fecundities according to their PGM-1 allozyme genotypes. Corrected fecundities by female size were determined on board from mature females by counting coelomic oocytes from aliquots and the *Pgm-1* genotype was determined in the laboratory. The numbers above each category indicate the number of individuals.

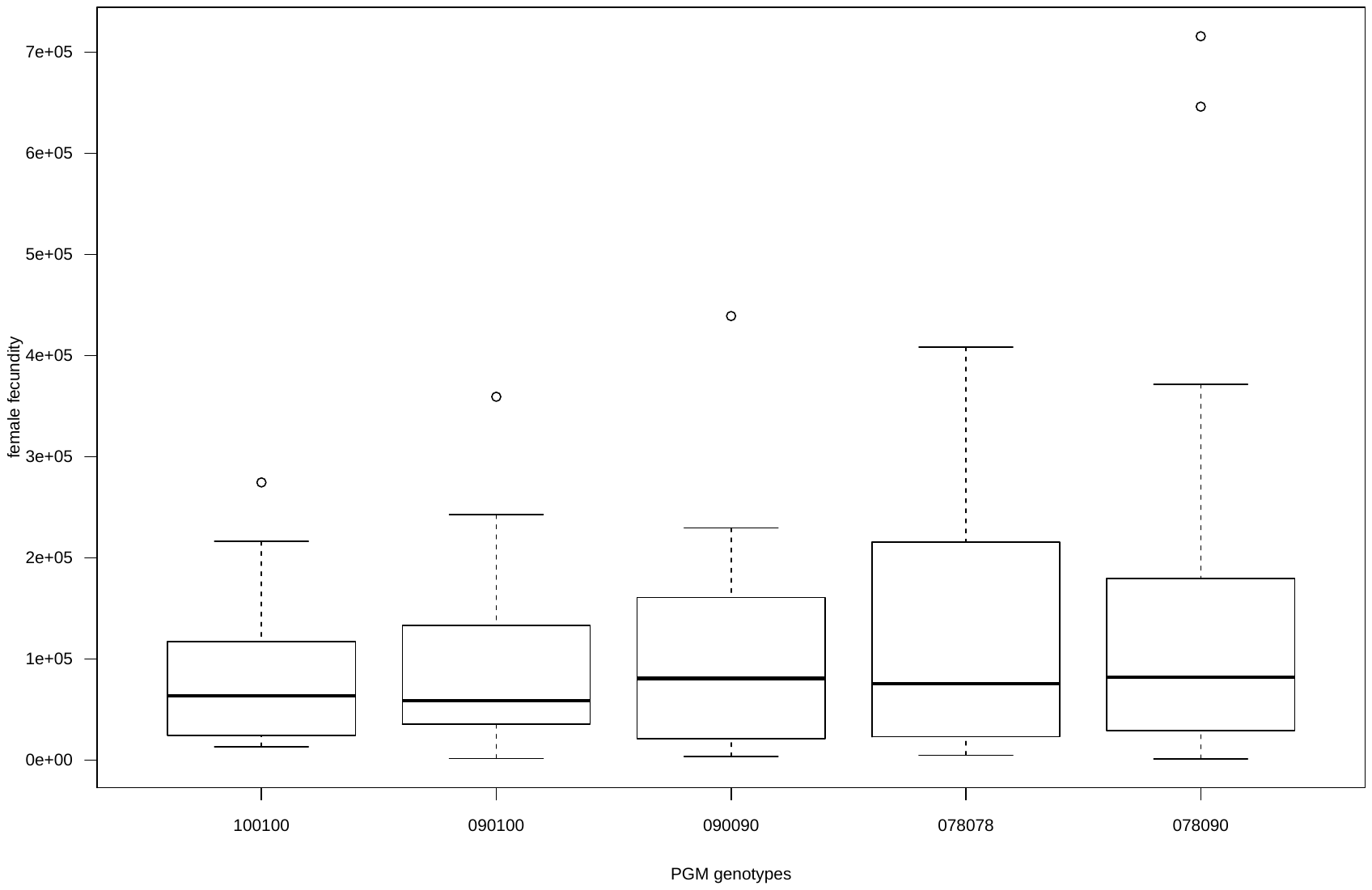
